# Exposure to immune stimuli reprograms alveolar macrophages to acquire neutrophil-derived antimicrobial molecules to prevent staphylococcal pneumonia

**DOI:** 10.64898/2025.12.03.692039

**Authors:** Amit Sharma, Linda M. Heffernan, Venkatesh Mayandi, Seetharama D. Jois, William N. Beavers, Basel H. Abuaita

## Abstract

The protective adaptation of innate immune defenses against respiratory pathogens has been linked to previous exposure to immune stimuli; however, the underlying mechanisms of these adaptations are not yet fully understood. Here, we show that pre-exposure to a low dose of non-specific immune stimuli or infection protects against subsequent lethal methicillin-resistant *Staphylococcus aureus* (MRSA) challenge. This enhanced protection concurs with increased alveolar macrophages (AMs) resulting from a self-renewal process in the lungs. Importantly, these AMs are programmed to acquire neutrophil-released antimicrobial enzymes from the extracellular space to kill MRSA, prevent tissue damage, and rapidly restore lung homeostasis. The gain of AM functions is dependent on differential gene expression, including expression of the efferocytosis receptor MerTK and the anti-apoptotic regulator Bcl-xL. Thus, our data highlight that the acquisition of neutrophil enzymes by AMs is an integral component of innate immune adaptation.

## INTRODUCTION

*Staphylococcus aureus* greatly impacts human health, causing nearly 25 million infections every year in the United States alone, and is the leading bacterial cause of death worldwide.^1,2^ *S. aureus* colonizes the skin and nasopharynx early in life, with approximately 70% of newborn babies becoming nasal positive for *S. aureus*.^3^ When opportunities are present, *S. aureus* is capable of infecting almost every niche of the body and causing diseases ranging from common superficial skin and soft tissue infections to more invasive and severe infections such as pneumonia, endocarditis, and sepsis.^4,5^ In addition, recurrent *S. aureus* infections result in serious problems, which frequently occur in more than 20% of infected patients.^6,7^ The molecular basis of the interindividual susceptibility to recurrent or invasive infections can be attributed to surgeries, intravenous devices, and disorders affecting the development and function of phagocytes.^8–12^. However, *S. aureus* infections also occur in otherwise healthy individuals, with some cases resulting in relapsing infections or progression to invasive disease. Yet, variations in immune responses among individuals has been observed and may confer predisposition to *S. aureus* infections.^13^ Individuals are constantly exposed to inflammatory stimuli such as bacterial endotoxin (lipopolysaccharides; LPS) and pathogens^14^, which can differentially reprogram host defenses and may account for the variations in patient susceptibility to *S. aureus*.

Effective immune responses against *S. aureus* involve cooperative interactions between innate immune cells, including neutrophils, monocytes, and macrophages.^15–17^ Once *S. aureus* invades tissue, resident macrophages and epithelial cells release immunomodulators to guide recruitment of blood circulating cells to the site of infection.^18,19^ Neutrophils are primary responder cells that migrate to the affected site within a few hours and kill large numbers of bacteria.^17^ Several mechanisms are utilized by neutrophils to kill extracellular bacteria including release of toxic enzymes, reactive oxygen species (ROS), and DNA into the extracellular matrix.^20^ These destructive molecules can also damage host tissues if not cleared in a spatiotemporal manner.^21–23^ Monocytes are also recruited to sites of infection and, along with resident macrophages, contribute to killing of bacteria as well as play a vital role in clearing damaged cells and toxic materials released into extracellular space to restore tissue hemostasis.^15,16,24,25^ Macrophages express surface molecules such as Tyro3, Axl, and Mer family receptors that interact with *eat-me* ligands on apoptotic cells to trigger their uptake through a process called efferocytosis.^26^ While the uptake of toxic materials from the extracellular space is critical for the resolution of inflammation, it is also possible that macrophages take these molecules up for future use. Recent studies have shown that primary innate immune cells isolated from various human patients differentially respond to immune stimuli.^27–29^ Although genetic variation greatly impacts these responses, non-heritable factors such as pre-exposure to microorganisms may also contribute to some of these variations. Certainly, repeated or prolonged exposure to low levels of inflammatory mediators can influence the function of innate immune cells.^30^ Nonetheless, whether pre-exposure to immune stimuli influences innate immune cells to prevent severe *S. aureus* infections remains unclear.

The lungs are constantly exposed to environmental and microbial particles that modulate inflammatory responses.^14^ Earlier studies showed that pre-exposure to immune stimuli, including LPS, Bacillus Calmette-Guérin (BCG) or microbes, induces resistance to subsequent bacterial lung infections.^31–35^ The protective responses gained during pre-exposure have been linked to modulation of AM and neutrophil phenotypes and functions. For example, initial exposure to immune stimuli modulates the AM surface marker phenotype and alters inflammatory responses to subsequent stimulation.^31–33^ Simultaneously, exposure to LPS or BCG modulates the neutrophil population and increases their expression of antimicrobial genes,^35,36^ In addition, it was demonstrated that mice exposed to LPS exhibit AMs resistant to cell death, which is often induced during subsequent immune challenges.^37^ These findings suggest that primary exposure to immune stimuli induces protective adaptations that involve cross-talk between AMs and neutrophils to better defend against pathogens. However, the underlying mechanisms of these adaptations are unknown.

Here, we used an *in vivo* model of MRSA-induced pneumonia to investigate the protective adaptations associated with repeated exposure to immune stimuli or pathogens. We found that prior stimulation with low doses of LPS, Pam3CSK4, Poly::IC, or non-lethal MRSA protects mice from a subsequent lethal MRSA infection. Pre-exposure to immune stimuli promotes AMs to uptake neutrophil-released antimicrobial agents to kill MRSA and prevent hyperinflammatory states. Our study thus deciphers an undescribed mechanism of macrophage host defense that occurs *in vivo,* where AMs acquire neutrophil payloads to promote MRSA clearance and resolution of inflammation.

## RESULTS

### Lung exposure to immune stimuli protects against subsequent MRSA pneumonia

Recurrent infection by *S. aureus* is common in humans; however, the mechanisms underlying susceptibility remain unclear. To establish a recurrent MRSA lung infection model in mice, we inoculated mice via the oropharyngeal route with a non-lethal dose of MRSA or control phosphate-buffered saline (PBS) for 6 days to introduce the bacteria into the lungs. Mice were then challenged with a lethal MRSA dose for 48 hours (Figure 1A). The 6-day timeline was chosen because it represents the time point at which primary MRSA infections are cleared and recruited neutrophils are resolved in the lungs (Figures 1B and 1C). The primary-infected mice exhibited severe signs of pneumonia, characterized by elevated inflammatory cells, dense clusters of bacteria, fibrin exudation, and necrosis (Figure S1A). As a result, 50% of mice died within 48 hours post-infection (hpi), and high bacterial counts were detected in the bronchoalveolar lavage fluid (BALF) and lungs (Figures 1D and 1E). By contrast, secondary-infected mice had a low grade of pneumonia, a 100% survival rate, and low bacterial burdens in the BALF and lungs. These data suggest pre-exposing mice to MRSA induces a protective pulmonary defense against secondary exposure.

**Figure 1.**
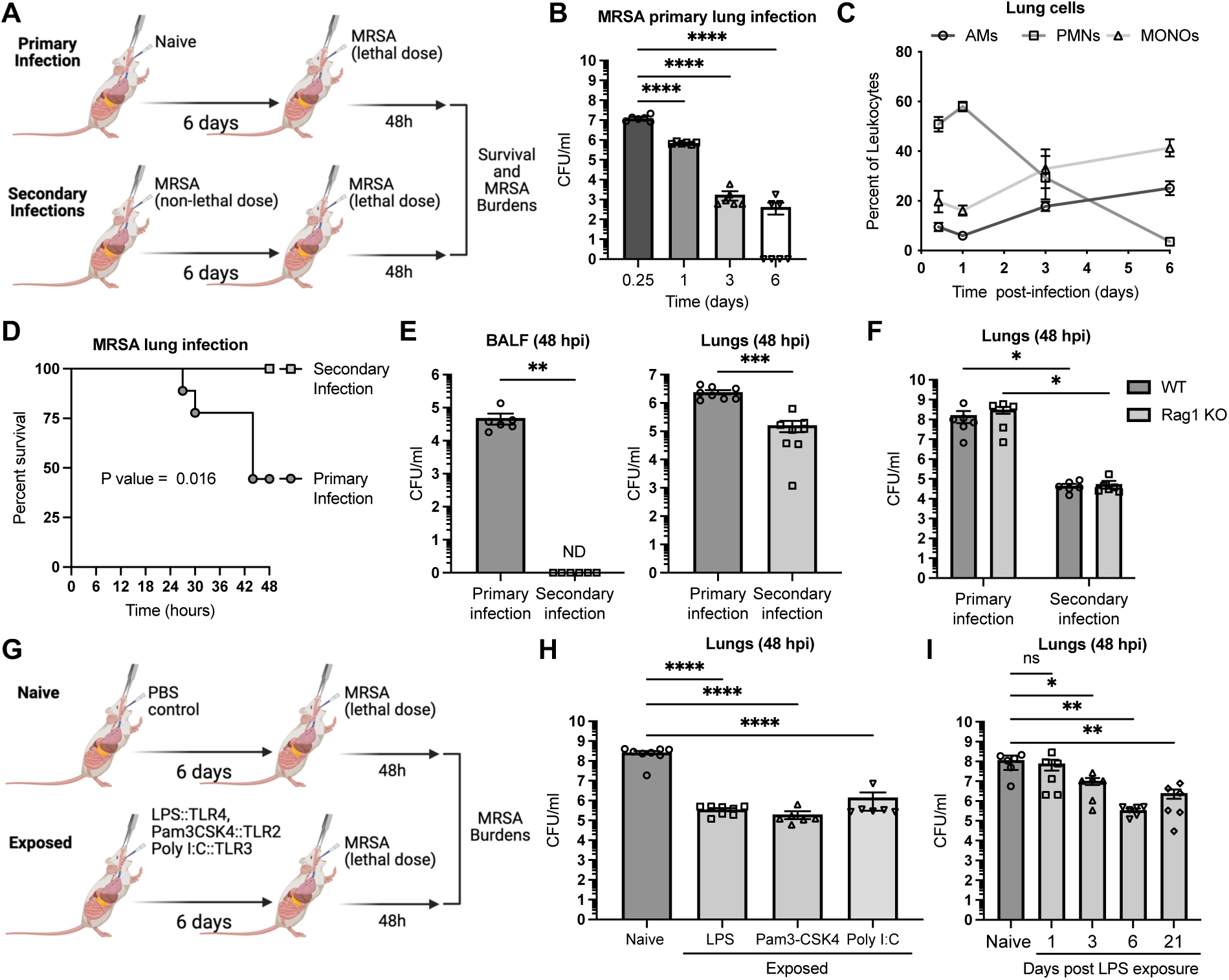
Lung exposure to non-lethal MRSA or TLR agonists protects against subsequent lethal MRSA infection. (A) Schematic showing primary and secondary (recurrent) MRSA infections. Mice were infected with lethal MRSA (primary) or initially infected with non-lethal MRSA before being rechallenged with lethal MRSA (secondary). (B) Time-course analysis of lung bacteria burdens of mice infected oropharyngel with MRSA (n≥6). (C) Time-course flow cytometry analysis of relative innate immune cells in lungs of MRSA-infected mice (n=4). Cells were stained with anti-CD11c/anti-CD170 for alveolar macrophages (AMs), anti-Ly6G/anti-CD11b for neutrophils (PMNs), and anti-Ly6G/anti-CD11b for monocytes (MONOs). (D) Survival curves of primary (n=9) and secondary (n=9) MRSA-infected mice. (E) Bacterial burdens in BALF and Lungs of primary (n≥6) and secondary (n≥6) MRSA-infected mice. (F) Bacterial burdens in lungs of WT or Rag1 KO mice infected primary (n≥6) and secondary (n≥6) with MRSA via oropharyngeal route. (G) Schematic representation of mice exposed to phosphate buffer saline (PBS; naive) or different TLR agonists (LPS, Pam3CSK4, or Poly::IC) before challenge with a lethal dose of MRSA. All inoculations were performed via the oropharyngeal route. (H) Bacterial burdens in lungs of naive mice (n=8) or mice exposed to LPS, Pam3CSK4, or Poly::IC for 6 days before MRSA lung infection (n=6). (I) Bacterial burdens in lungs of naive mice (n=6) or mice exposed to LPS for the indicated time before MRSA lung infection (n=6). Survival curves were graphed using a Kaplan-Meier survival plot, and the Log-rank (Mantel-Cox) test was used to calculate the P value. All other graphs represent the mean ± Standard Error of the Mean (SEM) from at least two independent experiments. *P* values for other graphs were calculated using Mann-Whitney test for panel E, One-way ANOVA followed by a Holm-Sidak multiple comparisons test for panels B, H, and I, and Two-way ANOVA followed by a Holm-Sidak multiple comparisons test for panel F. *P* value: *<0.05; **< 0.01, ***< 0.001, ****< 0.0001, and ns: not statistically significant.

To investigate whether innate or adaptive immune cells drive the protection against subsequent infection, we first challenged Rag1 knockout (KO) mice, which lack lymphocytes, with primary and secondary MRSA and assessed the bacterial burden at 48 hpi (Figure 1F). There were elevated and comparable MRSA counts in the lungs of both primary infected wild-type and Rag1 KO mice, indicating that lymphocytes do not contribute to the innate pulmonary immunity against primary MRSA infection. In addition, like wild-type mice, primary infection enhanced MRSA clearance during secondary infection in Rag1 KO mice, indicating that lymphocytes are also dispensable for the acquired protective immune response. Therefore, adaptive immune lymphocytes are unlikely to account for the observed protective response.

To assess whether protective immunity is part of a trained innate immune response acquired during exposure to non-specific immune stimuli, wild-type mice were exposed to different toll-like receptor (TLR) agonists followed by challenge with a lethal dose of MRSA. Mice exposed to TLR-2, 3, or 4 agonists were protected against MRSA infection and exhibited low MRSA counts in the lungs compared to naive mice (Figures 1G and 1H). Importantly, exposure of mice to immune stimuli for one day was insufficient to protect against MRSA (Figure 1I), indicating that the initial immune activation requires some time to develop. However, once acquired protective immunity is induced, it lasts for at least 21 days (Figure 1I). Thus, our data suggest that exposure to immune stimulation, such as MRSA or Toll-like receptors (TLR) agonists, leads to protective immune responses against subsequent MRSA challenges, a hallmark of trained immunity.

### Neutrophils are essential for LPS-induced pulmonary immunity against MRSA

Recently, we established that innate immune cells, including alveolar macrophages (AMs), inflammatory monocytes (MONOs), and neutrophils, also known as polymorphonuclear leukocytes (PMNs) collaboratively promote a protective pulmonary response against MRSA challenge.^16^ To profile the dynamics of these cells in the lungs upon immune stimulation, we performed cellular immunophenotyping in the lungs and BALF of mice exposed to LPS for 3 or 6 days (Figures 2A and S2A). Similar to MRSA primary infection, neutrophil infiltration in the lungs was observed on day 3 and then resolved by day 6. Monocyte infiltration was also increased on day 3 but remained constant through day 6. By contrast, the relative level of AMs was decreased on day 3 and then replenished by day 6. These data suggest that changes in the composition of innate immune cell populations following LPS exposure may drive the observed protective response against MRSA challenge.

**Figure 2.**
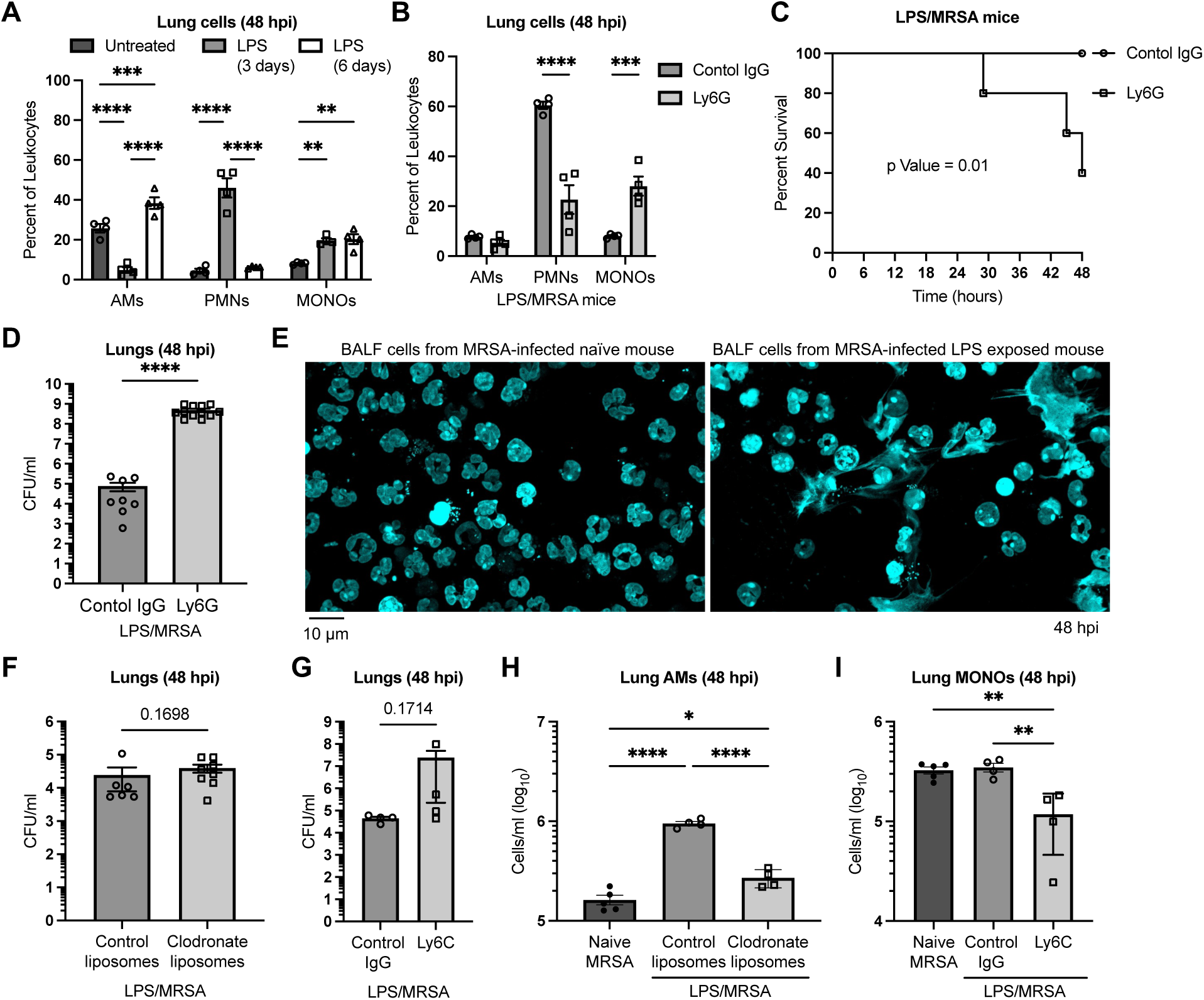
Neutrophils and alveolar macrophages are required for the acquired protective adaptations during LPS exposure against MRSA lethal challenges. (A) Immunophenotyping analysis of leukocytes in lungs of naive mice (n=4) and mice exposed to LPS via oropharyngeal aspiration for 3 or 6 days (n=4). Cells were stained for AM markers (anti-CD11c, anti-CD170), neutrophil (PMN) markers (anti-Ly6G, anti-CD11b), and monocyte (MONO) markers (anti-Ly6C, anti-CD11b) and analyzed by flow cytometry. (B) Immunophenotyping analysis of leukocytes in lungs of LPS-exposed mice when injected with control IgG (n=4) or anti-Ly6G (n=4) intravenously for two consecutive days before MRSA lung infection. Cells were stained as in panel A. (C, D) Survival (C) and bacterial burden (D) of LPS-exposed mice when mice were injected with control IgG (n≥8) or anti-Ly6G (n≥8) intravenously for two consecutive days before MRSA lung infection. (E) Representative confocal microscopy images of BALF cells from MRSA-infected naive and LPS-exposed mice. Cells were stained with Hoechst 33342 (Gray) to label DNA. (F, G) Bacterial burden (F) and alveolar macrophage count (G) in lungs of LPS-exposed mice for 6 days, administered control liposomes (n≥4) or clodronate liposomes (n≥4) in two consecutive days, and subsequently challenged with MRSA. (H, I) Bacterial burden (H) and monocyte count (I) in lungs of LPS-exposed mice for 6 days before MRSA infection. Mice were injected with control IgG (n≥4) or anti-Ly6G (n≥4) intravenously for two consecutive days prior to LPS exposure. All graphs represent data from at least two independent experiments. Survival curves were graphed using a Kaplan-Meier survival plot, and the p-value was calculated by the Log-rank (Mantel-Cox) test. All other graphs represent the mean ± SEM and the p values were calculated by Mann-Whitney test for panel D, F, and G, One-way ANOVA followed by a Holm-Sidak multiple comparisons test for panels H, and I, and Two-way ANOVA followed by a Holm-Sidak multiple comparisons test for panels A and B. *P* value: *P* value: *<0.05; **< 0.01, ***< 0.001, ****< 0.0001, or exact numeric value.

To define the contribution of AMs, PMNs, and MONOs in acquired immunity, these cells were singly depleted and the susceptibility of LPS-exposed mice to subsequent MRSA challenges was assessed. PMNs and MONOs were depleted by administering anti-ly6G or anti-ly6C, respectively, in the blood via tail vein injection. AMs were depleted by inoculating mice with clodronate liposomes via the oropharyngeal route as previously performed.^16^ Depletion of PMNs rendered mice highly susceptible to MRSA challenge as evidenced by low PMN counts and high MRSA burdens in the lung, which resulted in a poor survival rate (Figures 2B-2D). Remarkably, we readily observed more released neutrophil extracellular traps (NETs) in the BALF of LPS-exposed mice when compared to naive mice (Figure 2E). These data suggest that PMNs and NET release may play a role in LPS-induced protective immunity. However, adoptive transfer of PMNs isolated from LPS-exposed mice to naive mice failed to induce protection against MRSA challenge (Figures 2SB and 2SC). Thus, LPS exposure may induce localized environmental cues in the lungs that are required for PMN-dependent immunity. In contrast to PMNs, depletion of AMs and MONOs exhibited no measurable effect on bacterial burdens in the lungs of LPS-exposed mice despite the reduction of absolute cell counts (Figures 2F-2I). Although AM numbers were reduced in clodronate-inoculated LPS-exposed mice relative to controls, absolute counts remained largely higher than in naive mice (Figure 2H). This may indicate that the remaining AMs were resilient to depletion and were sufficient to protect against MRSA challenge.

As an additional way to assess the role of inflammatory monocytes in LPS-induced immunity, we used CCR2 KO mice, which are defective in MONO recruitment to inflamed sites.^38^ Exposure of CCR2 KO mice to LPS resulted in a similar level of protection as seen in wild-type mice despite the reduction of MONO counts (Figures S2D and S2E). In addition, we observed a higher number of AMs in the lungs of LPS-exposed WT and CCR2 KO mice during MRSA infection compared to naive WT-infected mice, despite the reduction in MONOs in CCR2 KO mice (Figure S2F and S2G). This may imply that the increase in AM counts in infected lungs of LPS-exposed mice occurs independently of CCR2-mediated monocyte recruitment. Collectively, our data suggest that both PMNs and AMs contribute to the immune protection induced by LPS exposure.

### LPS exposure increases the initiation and resolution of inflammation during subsequent MRSA challenges

To explore the role of pre-exposure to immune stimuli in the kinetic activation and resolution of pulmonary immune responses, we first monitored the transcript levels of inflammatory mediators at 6 hours post-infection (6 hpi), thus during the initiation period of MRSA pneumonia. We found that most inflammatory mediators were highly induced in the lungs of mice when pre-exposed to LPS relative to naive mice (Figure 3A). It is worth noting that there were low levels of inflammatory genes in the lungs of LPS-exposed mice before MRSA infection (Figure S3A). Thus, the increase in the transcription response in the lungs of LPS-exposed mice during early MRSA infection was not due to residual transcripts from the pre-exposure. We next used NanoString technology to quantify 785 transcripts of mouse inflammatory response genes to get a more detailed understanding of the protective pulmonary host response induced during LPS exposure. The number of inflammatory genes with expression that changed significantly in response to MRSA was higher in the lungs of LPS-exposed mice than in the lungs of naive mice (Figure 3B). This was true for the number of upregulated genes, but not downregulated genes. Interestingly, we found that some genes were inversely changed during MRSA challenge in the lungs of LPS-exposed mice relative to naive mice. These genes encode the scavenger receptor (Marco), Alanine aminopeptidase (Anpep), and the complement receptor 4 subunit (Itgax), which were increased in MRSA-infected lungs of LPS-exposed mice and decreased in naive mice (Figure S3B). To further characterize other genes that changed in MRSA-infected mice in response to LPS pre-exposure, we analyzed genes with differentially changed expression in LPS-exposed mice compared to naive mice during MRSA infection and identified 406 genes (Figure 3C and Table S1). This gene set was further filtered to include genes that were not significantly changed in MRSA-infected naive mice, representing unique genes altered only during MRSA infection in LPS-exposed mice (Figure S3C and Table S2). The list of unique genes contained antimicrobial molecules (Nos2, Gbp2, Gbp2, and Lyz2), lysosomal degradative enzymes (Psap, Ctsl, Ctss, Ctsw, Ctsz, and Gusb), phagocytic molecules (C3ar1, Cd68, Marco, Itgb2, and Itgax), and regulators of programmed cell death (Fas, Bcl2l1, Xaf, Bax, and Ddit3). These data imply that genes uniquely differentially expressed in LPS-exposed mice may contribute to the protective immune response against subsequent MRSA challenge.

**Figure 3.**
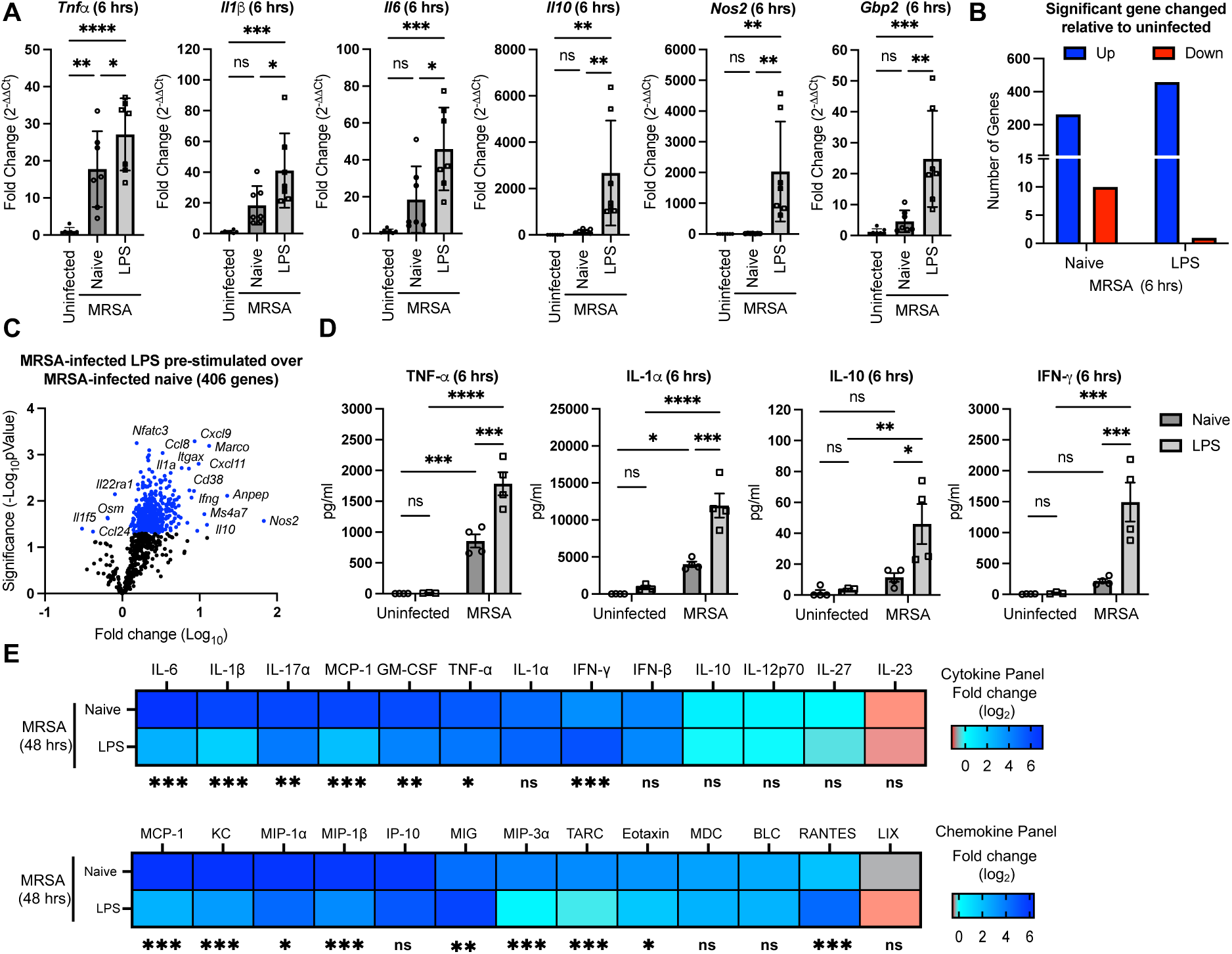
LPS exposure modulates the magnitude of inflammatory responses during the early and late stages of MRSA infection. (A) Expression of indicative inflammatory genes in lungs of uninfected mice, MRSA-infected naive mice, and MRSA-infected LPS pre-exposed mice was assessed by quantitative real-time PCR (qPCR) relative to *Gapdh*. The ΔΔCt was calculated relative to the average ΔCt of uninfected mice. (B) Number of significantly changed genes in MRSA-infected lungs of naive or LPS-pre-exposed mice relative to uninfected mice. Upregulated (Blue) or downregulated (Red) genes were identified based on a *P* value < 0.05 of the log_10_ ratio of infected over uninfected normalized counts. (C) Volcano plot of differentially expressed genes in MRSA-infected lungs from LPS pre-exposed mice over MRSA-infected lungs from naive mice. Blue dots indicate genes that were significantly changed. (D) Levels of inflammatory cytokines produced in the lungs of uninfected and MRSA-infected mice for 6 hours when mice were pre-exposed to LPS for 6 days or left unexposed (naive) mice. Cytokine levels in acellular fractions of lung single cell suspensions were quantified by flow cytometry using a LEGENDplex bead-based assay. (E) Expression of cytokines and chemokines in lungs of MRSA-infected mice at 48 hpi when mice were left unexposed (naive) or pre-exposed to LPS for 6 days. Heatmaps representing the mean of fold change of infected mice (n=4) relative to uninfected mice (n=4). Other graphs represent the mean of n≥3 biological replicates ± standard deviation (SD). *P* values were calculated using One-way ANOVA for panel A or Two-way ANOVA for panels D and E, followed by a Holm-Sidak’s multiple comparisons test. *P* value: *<0.05; **< 0.01, ***< 0.001, ****< 0.0001, and ns: not statistically significant.

We next sought to determine whether the initially induced transcriptional responses during MRSA infection occur at the protein level and whether they change during the resolution stage of pneumonia. A bead-based multiplex assay was used to quantify the level of cytokines produced in the lungs of MRSA-infected mice at 6 and 48 hours. Consistent with the transcriptional responses, LPS-exposed mice exhibited increased levels of several cytokines (IL-1α, IL-17α, IL-10, IFN-γ, and IFN-β) during MRSA infection relative to MRSA-infected naive mice (Figures 3D and S3D). Yet, some other pro-inflammatory cytokines, such as IL-1β, IL-6, IL-12, IL-23, IL-27, and GM-CSF, were present at lower levels in MRSA-infected LPS-exposed mice compared to naive mice (Figure S3D). Thus, the initial heightened inflammatory response only encompasses a select group of proteins. Compared to the initial stage of inflammation, the levels of most quantified cytokines and chemokines, except for IFN-γ, MIG and Rantes, were produced in substantially lower levels at the late period of infection (48 hpi) in LPS-exposed mice relative to naive mice during MRSA infection. This profile of rapid onset and quick resolution of inflammation that we observed is similar to the inflammatory response profile observed during trained immunity.^39^ Taken together, these data demonstrate that exposure to immune stimuli enhances the initiation and resolution of immune responses, thereby protecting against MRSA-induced pneumonia.

### Exposure to immune stimuli reprograms self-renewed AMs to scavenge PMN-derived antimicrobial enzymes

Visualizing innate immune cells in infected organs can lead to unexpected insights. Thus, microscopy was used to visualize innate immune cells from the lungs of LPS-exposed and naive mice during MRSA infection. Large cell populations were readily observed in BALF of LPS-exposed mice that often-contained double nuclei, suggesting that they were dividing (Figure S4A). In addition, most of these large cell populations in the lungs were double-positive for CD11c and CD169 and were highly phagocytic, as there was a large presence of intracellular MRSA (Figures 4A and S4B). These observations suggest that the large, highly phagocytic cells are AMs undergoing cell expansion in lungs of LPS-exposed mice during infection. Concurring with the observations, we found that there were higher AM counts in MRSA-infected lungs of LPS-exposed mice when compared to naive mice (Figure S4C).

**Figure 4.**
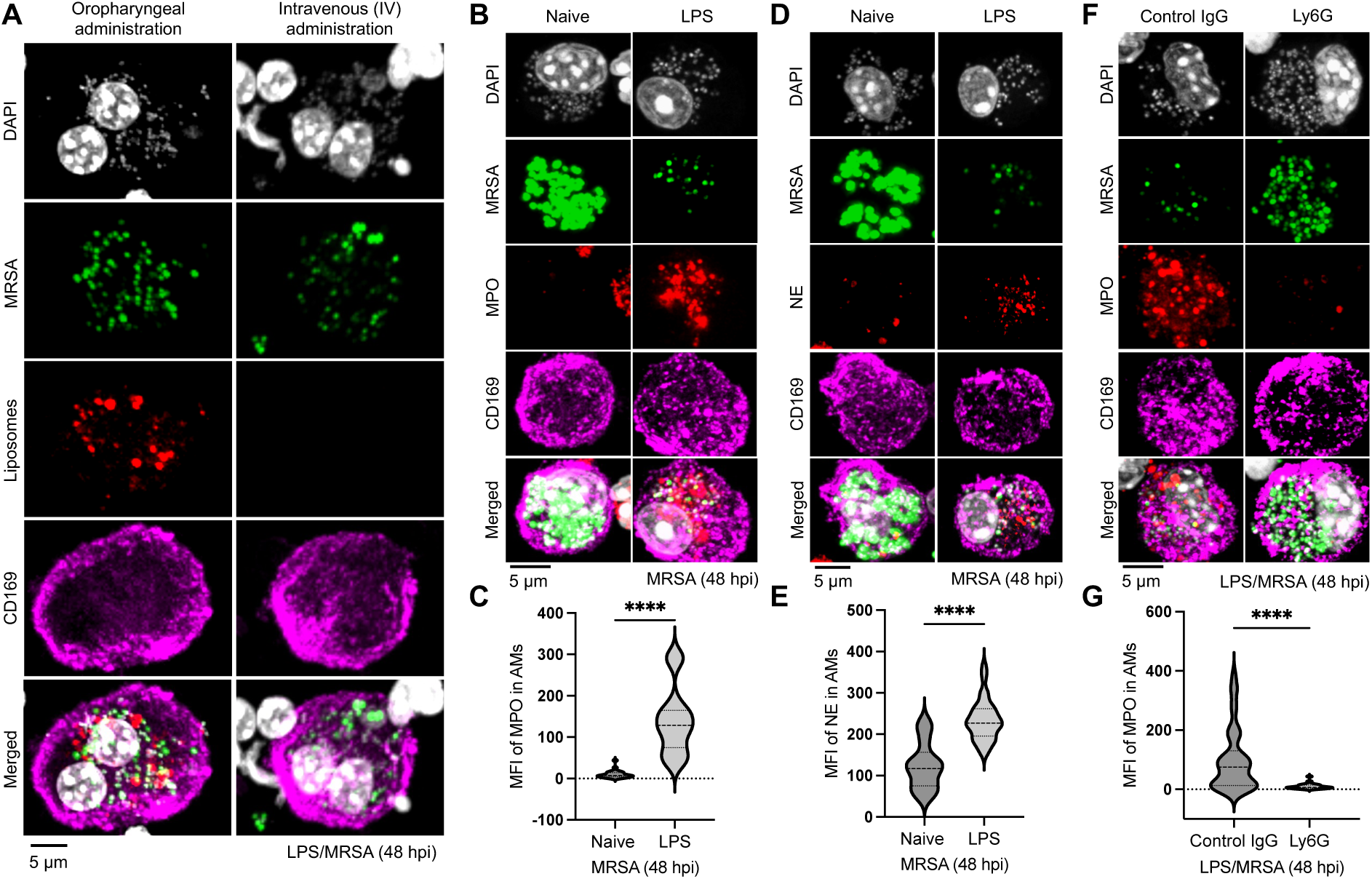
Resident AMs undergo self-renewal and reprogramming to uptake PMN-derived antimicrobial enzymes during LPS exposure. (A) Representative confocal microscopy images of resident AMs labeled with Rhodamine liposomes (red) before mice were exposed to LPS for 6 days and subsequently infected with MRSA-GFP (green). Binucleated AMs were identified based on CD169 positive staining (magenta) and presence of double nucleus based on DAPI staining (gray). (B, C) Representative confocal microscopy images (B) and quantification (C) of MPO levels in AMs from MRSA-GFP (green) infected lungs of unexposed (naive) mice or LPS pre-exposed mice for 6 days. Cells were collected *ex vivo* from lungs, adhered to coverslips, and stained with DAPI (gray), anti-MPO (red), and CD169 (magenta). (D, E) Representative confocal microscopy images (B) and quantification (C) of neutrophil elastase (NE) levels in AMs when naive or LPS pre-exposed mice were infected with MRSA-GFP (green). Cells were extracted from the lungs and stained with DAPI (gray), anti-NE (red), and CD169 (magenta). (F, G) Representative confocal microscopy images (B) and quantification (C) of MPO levels in AMs of LPS pre-exposed mice when mice were injected twice via tail vein with control IgG or anti-Ly6G and subsequently infected with MRSA-GFP (green) for 48 hours. Lung cells were extracted *ex vivo* and stained with DAPI (gray), anti-MPO (red), and CD169 (magenta). Mean fluorescence intensity (MFI) of MPO or NE levels in AMs was quantified by ImageJ software. Graphs are represented as violin plots of MFIs of at least 60 cells for each condition from n≥3 mice. *P* values were calculated using Mann-Whitney tests. *P* value: *< 0.05, ***<0.001, ****<0.0001. Scale bars represent 5 µm.

To investigate the origin of the expanded AMs, we assessed whether these cells originate from resident AMs in BALF or from recruited MONOs from blood. Mice were inoculated with fluorescent (Rhodamine B) liposomes oropharyngeally to label resident AMs in BALF or intravenously to label blood MONOs one day before LPS exposure. Mice were then challenged with MRSA-GFP at day 6 post-LPS exposure and BALF cells were visualized at 48 hpi by confocal microscopy. The administration of fluorescent liposomes in BALF was traced into the highly phagocytic double-nuclei AMs, indicating that these cells originate from resident AMs (Figure 4A). In contrast, intravenous injection of fluorescent liposomes did not trace into the highly phagocytic double-nuclei AMs, despite effective blood MONO labeling (Figures 4A and S4D). These data suggest that the increased AM number in the lungs of LPS-exposed mice is a result of self-renewing resistant AMs, and these cells are highly phagocytic.

To characterize the cooperative host defenses between macrophages and neutrophils in infected lungs, single-cell suspensions were extracted, stained, and visualized by confocal microscopy from lungs from mice infected with MRSA-GFP. Macrophages were labeled with anti-CD169, and neutrophils were labeled with either anti-Myeloperoxidase (MPO) or anti-Neutrophil Elastase (NE). We first observed that the large macrophages from LPS-exposed mice contained lower fluorescence GFP bacteria when compared to macrophages from naive mice (Figure 4B). This observation was confirmed when the MRSA-GFP fluorescence intensity of infected lung cells was quantified by flow cytometry (Figure S4E), suggesting that the reduction in MRSA-GFP fluorescence intensity could be due to killed MRSA. Unexpectedly, we found that the large macrophages from LPS-exposed lungs were also positive for neutrophil markers (MPO and NE) (Figures 4B-4E). To explore whether MPO was induced in macrophages by exposing mice to LPS or was acquired from extracellular environments, we first quantified *Mpo* transcripts from LPS-exposed and naive mice during MRSA infection. LPS-exposed mice had lower levels of *Mpo* transcript relative to naive mice during MRSA challenge (Figure S4F), suggesting that an increase in *Mpo* expression is not likely the mechanism underlying the accumulation of MPO in macrophages. Similarly, LPS-exposed mice exhibited lower levels of neutrophil elastase (*Elane*) expression in response to MRSA challenges compared to naive mice (Figure S4G). We then monitored the kinetics of MPO accumulation in macrophages at different time points post-MRSA challenge. There were low MPO-positive macrophages in lungs at an earlier time point (6 and 24 hpi) and high MPO-positive AMs at a later time point (48 hpi) (Figure S4H), indicating the MPO gradually accumulated in macrophages over time. In addition, we observed different sizes of extracellular MPO-positive vesicles in lungs at an earlier time point (24 hrs), which implies that the MPO and NE were taken up from the extracellular space (Figure S4J). Importantly, neutrophil depletion significantly reduced the number of MPO-positive AMs (Figure S4C). These observations suggest that neutrophil MPO and NE were released into the lung extracellular space during the initial stage of infection and then were subsequently acquired by AMs.

### MerTK is required for MPO uptake by AMs to prevent lung injuries

PMNs release various substances to combat pathogens; however, the clearance of these molecules via a process called efferocytosis is also essential to prevent damage to the surrounding tissues.^25,40^ A previous study suggested that macrophages can capture PMN-released antimicrobial substances and exploit them to kill microbes.^41^ We readily observed MPO localized with MRSA and Lamp1 (Figure 5A), indicating MPO is sequestered into phagosomes. Thus, we hypothesized that AMs take up MPO to kill MRSA and limit lung injuries. To test this hypothesis, we explored whether MPO inhibition weakened the protective immunity acquired during LPS exposure. Mice were exposed to LPS via the oropharyngeal route for 6 days and then treated with MPO inhibitor (4-Aminobenzoic acid hydrazide; 4-ABAH) or vehicle control 24 h before MRSA lung infection and then every 24 h throughout the duration of the experiment. Mice treated with 4-ABAH exhibited a 100% survival rate and similar lung bacterial clearance when compared to vehicle control (Figure 5B), indicating that MPO inhibition is insufficient to mitigate the protective immunity acquired during LPS exposure.

**Figure 5.**
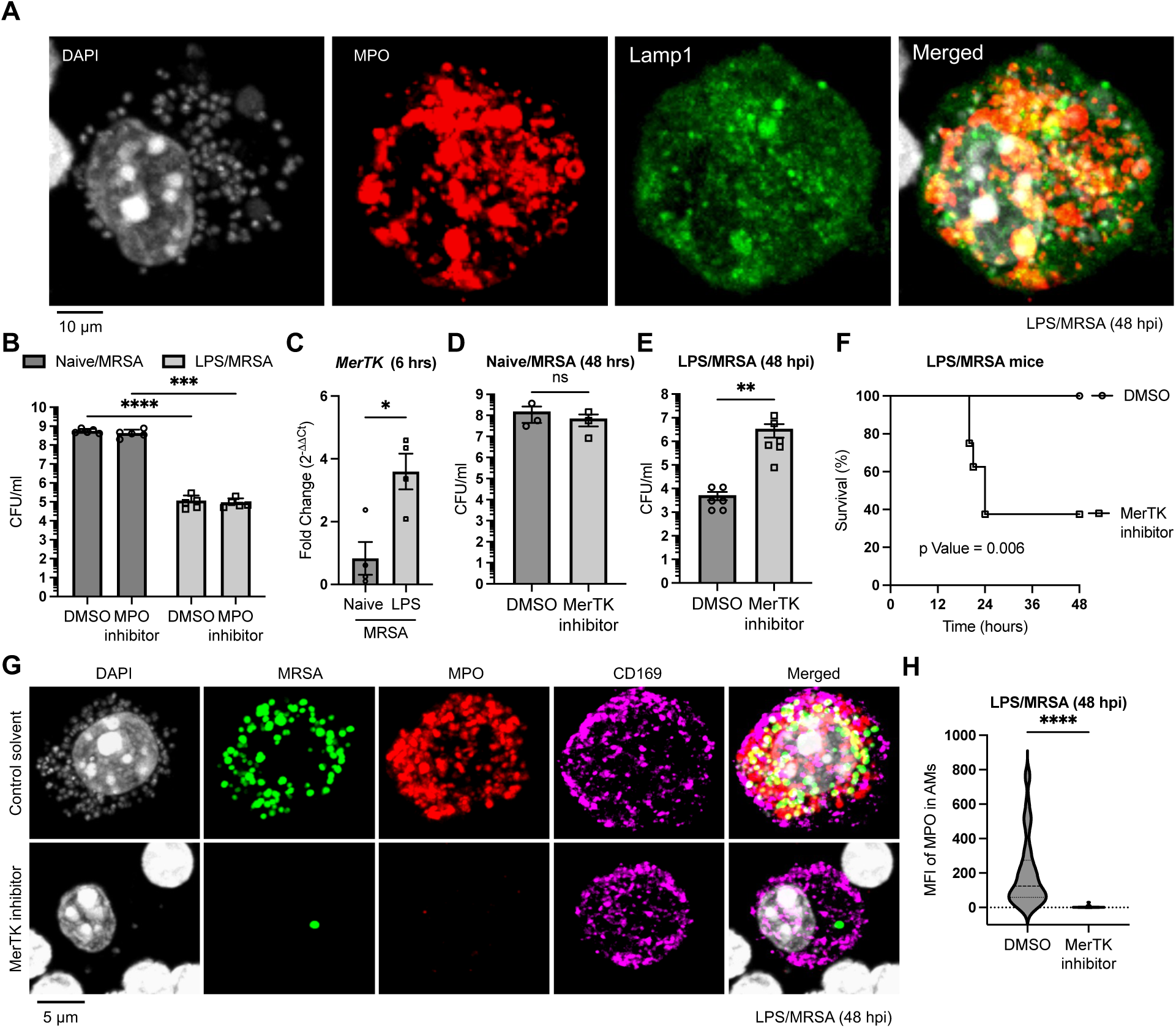
MerTK-mediated phagocytosis is essential for LPS-induced immunity against MRSA. (A) Representative confocal microscopy images of AMs collected from lungs of mice pre-exposed to LPS for 6 days and infected with MRSA for 48 hours. Cells were stained with DAPI (gray), anti-MPO (red), and anti-Lamp1 (green). (B) Bacterial burden in lungs of MRSA-infected naive mice treated with DMSO control (n=5), MRSA-infected naive mice treated with MPO inhibitor (n=5), MRSA-infected mice pre-exposed to LPS and treated with DMSO control (n=5), and MRSA-infected mice pre-exposed to LPS and treated with MPO inhibitor (n=5). (C) The expression level of *MertK* was quantified by qPCR and normalized relative to *Gapdh*. The ΔΔCt was calculated relative to the average ΔCt of uninfected mice. (D) Bacterial burden in lungs of MRSA-infected mice when mice (n=6) were treated with MertK inhibitor (n=6) or DMSO control (n=7). (E, F) Bacterial burden (E) and survival (F) of LPS-exposed mice when mice were treated with DMSO control (n≥6) or MertK inhibitor (n≥6) and subsequently infected with MRSA. (G, H) Representative confocal microscopy images (G) and quantification (H) of MPO accumulation in AMs extracted from lungs of LPS-exposed mice for 6 days, treated with DMSO control or MerTK inhibitor prior to being infected with MRSA-GFP (green) for 48 hours. Cells were stained with DAPI (gray), anti-MPO (red), and anti-CD169 (magenta. Violin plots represent MPO MPI of at least 60 cells for each condition from n≥3 mice. Other graphs represent the mean ± SEM from at least two independent experiments. *P* values were calculated using Two-way ANOVA followed by a Holm-Sidak’s multiple comparisons test for panel B, Mann-Whitney test for panels C, D, E, and H, or Log-rank (Mantel-Cox) test for panel F. *P* value: *<0.05; **< 0.01, ***< 0.001, ****< 0.0001, and ns: not statistically significant.

We next tested whether clearing damaging neutrophil-derived molecules, including MPO via efferocytosis, is required for the observed MRSA protection in LPS-exposed mice. Previous studies demonstrated that Mer proto-oncogene tyrosine kinase (MerTK) plays a vital role in the clearance of apoptotic cells and MPO.^42,43^ In addition, we observed that the expression of MerTK is increased in MRSA-infected lungs of mice exposed to LPS (Figure 5C), anticipating that MerTK is a suitable candidate for MPO clearance from MRSA-infected lungs. Thus, we assessed whether MerTK inhibition interferes with the protective immunity against MRSA challenge in LPS-exposed mice. MerTK inhibitor (UNC2025) neither induced lung cell death of uninfected mice nor interfered with MRSA clearance from lungs of MRSA-infected naive mice (Figures S5D and S5A). However, treating LPS-exposed mice with MerTK inhibitor during MRSA infection increased bacterial burdens in the lungs, lowered the survival rate of mice, and reduced MPO uptake by AMs (Figures 5E-5H). These findings suggest that MerTK is essential in MPO clearance from MRSA-infected lungs, which is part of the protective immune response acquired during LPS pre-stimulation.

### Bcl-xL is essential for AM-mediated MPO clearance in response to LPS exposure

A previous study suggested that AMs become more resilient to cell death during repeated exposure to inflammatory stimuli.^37^ We observed that AM depletion using chlodronate, which induces intrinsic apoptosis, was less effective in mice exposed to LPS in contrast to our previous study in naive mice (Figure 2H).^16^ Thus, we hypothesized that immune stimulation primes AMs to become more resilient to cell death by upregulating anti-apoptotic genes during subsequent infections. To test this hypothesis, we first evaluated changes in apoptosis-related genes from the NanoString transcriptional profiles of MRSA-infected mice when exposed to LPS. Expression of several anti-apoptotic genes, including *Bcl-2l1, Mcl1 and Bcl2,* was increased in MRSA-infected lungs of mice exposed to LPS relative to naive mice (Figure 6A). In contrast, expression of pro-apoptotic genes, such as *Bax and Bnip3*, remained unchanged. To evaluate the requirement of anti-apoptotic gene activity in immunity against MRSA, we used small molecule inhibitors to selectively target Bcl-2 (Venetoclax) or Bcl-xL (A-1331852). Mice treated with inhibitors via oropharyngeal inoculation did not exhibit lung toxicity as assessed by H&E-stained histology sections (Figure S6A). Remarkably, LPS-exposed mice treated with the Bcl-xL inhibitor were significantly susceptible to subsequent MRSA challenges, while those treated with the Bcl-2 inhibitor were not. This was evidenced by the higher mortality rates and bacterial burdens in the lungs when compared to control solvent (Figures 6B and 6C). The increased susceptibility of Bcl-xL inhibitor-treated mice was not due to interference with immune cell recruitment, as similar relative numbers of AMs, PMNs, and MONOs were present when compared to control solvent treatment (Figure 6D). Furthermore, confocal microscopic analysis showed less MPO accumulation in AMs of mice treated with the inhibitors (Figures 6E and 6F). Surprisingly, AMs from mice treated with inhibitors were negative for cleaved caspase 3 (Figure S5C), suggesting that Bcl-xL may contribute to AM function independently of protecting the cells from infection-induced cell death. However, other cell types from the lungs of MRSA-infected mice treated with Bcl-xL and MerTK inhibitors were positive for cleaved caspase 3. The death of these cells may contribute to the loss of protection against MRSA challenge when mice were treated with inhibitors. Collectively, Bcl-xL is upregulated in infected lungs of mice that have been exposed to LPS, which is essential for MPO uptake by AMs and immunity against MRSA challenge.

**Figure 6.**
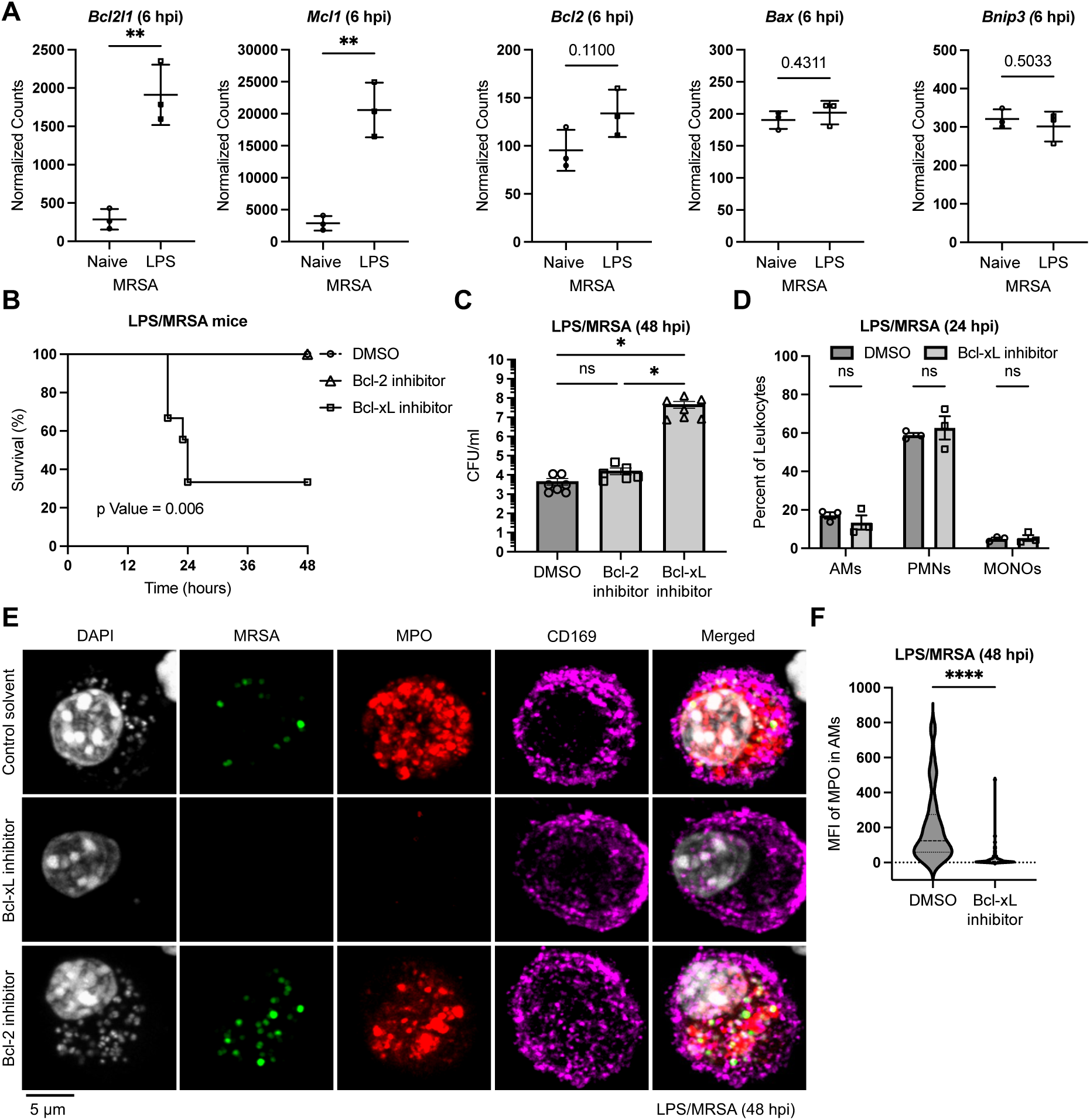
Induction of Bcl-xL in response to LPS exposure is required for acquired innate immunity against MRSA infection. (A) Normalized transcript levels of apoptosis-related genes (*Bcl2l1*, *Mcl1*, *Bcl2*, *Bax*, and *Bnip3*) in lungs of MRSA-infected naive mice (n=3) or LPS pre-exposed mice (n=3) from NanoString data. (B, C) Survival (B) and bacterial burden (C) of LPS pre-exposed mice for 6 days, treated with DMSO control (n≥7), Bcl-2 inhibitor (Venetoclax; n≥6), or Bcl-xL inhibitor (A-1331852; n≥7), and subsequently infected with MRSA. (D) Immunophenotyping analysis of leukocytes in lungs of MRSA-infected mice when injected with DMSO (n=3) or Bcl-xL inhibitor (n=7). (E, F) Representative confocal microscopy images (E) and quantification (F) of MPO accumulation in AMs from lungs of LPS-exposed mice for 6 days, treated with DMSO control, Bcl-2 inhibitor, or Bcl-xL inhibitor prior to being infected with MRSA-GFP (green) for 48 hours. Cells were stained with DAPI (gray), anti-MPO (red), and anti-CD169 (magenta). Violin plots indicate MFI of MPO from at least 60 cells for each condition from n≥3 mice. All other graphs indicate the mean ± SEM from at least two independent experiments. *P* values were calculated using Mann-Whitney test for panel A, Log-rank (Mantel-Cox) test for panel B, One-way ANOVA followed by a Holm-Sidak’s multiple comparisons test for panel C, or Two-way ANOVA followed by a Holm-Sidak’s multiple comparisons test for panel D. *P* value: *<0.05; **< 0.01, ***< 0.001, ****< 0.0001, ns: not statistically significant, or exact numeric value.

## DISCUSSION

Humans are constantly exposed to various inflammatory stimuli both externally from the environment and internally during recurrent infections.^14^ However, how previous exposure shapes inflammatory responses to subsequent infection remains unclear. We report that mice exposed to a non-lethal dose of LPS, Pam3CSK4, Poly:: IC, or MRSA via the oropharyngeal route exhibited increased survival against subsequent lethal MRSA challenge. This pre-exposure increases MRSA clearance and drives rapid resolution of inflammation when compared to naïve mice. The enhanced protective response correlates with an increase in self-renewal of alveolar macrophages (AMs), as evidenced by the presence of double-nuclei proliferating cells which traced back to resident AMs. Importantly, AMs were programmed to uptake neutrophil antimicrobial enzymes from the matrix to kill MRSA, prevent tissue damage, and rapidly restore lung hemostasis. Specifically, AMs induce expression of unique genes, including MerTK and Bcl-xL, to drive protective immune adaptations against MRSA via increasing efferocytosis. Collectively, our data highlight the importance of self-renewed AMs and enhanced acquisition of neutrophil enzymes by efferocytosis in the protective innate immune adaptations gained in response to pre-exposure to immune stimuli.

The concept of trained immunity has recently been described as an innate immune adaptation of cells that was gained during recurrent exposure to immune stimuli after cells returned to a non-activated state.^39^ Nevertheless, whether the cells must return to a normal state to gain the trained immune adaptations has not been demonstrated. Return of cells to the normal state is the discrimination between the signal from the initial inflammation and the second trained innate immune responses. In addition, the difference between activated and non-activated states is based on selective inflammatory genes. In our model system, we found that the initial inflammation must decline to induce the protective response against MRSA. We found that when the initial inflammation remained high at 24 h post-LPS exposure, mice were not protected against MRSA. However, mice were protected against MRSA when exposed to LPS for at least 3 days, even if there was residual inflammation. The timeline for this protection of at least 3 days after LPS exposure and lasting for several weeks is also aligned with other recent reports.^31,32,35^ Thus, we speculate that trained immunity is induced early on during exposure to immune stimuli, but a resolution period must be conceded to distinguish between the initial and trained immune responses.

Exposure to inflammatory stimuli induces AMs to undergo expansion and phenotypic changes. Our AM quantification was performed based on the counts of double-positive CD11c and CD170 cells. We found that the absolute count of AMs increased in mice exposed to LPS when compared to naive mice. Similar findings were also reported in a previous study where AM counts decreased when mice were exposed to LPS and then rebounded by day 6.^37^ A more recent study suggests that AMs undergo phenotypic plasticity in response to immune stimuli rather than cell expansion.^31^ Specifically, it was shown that mice exposed to LPS for 14 days have 4 clusters of AMs based on surface markers in their lungs. These AM subsets were described as original *Siglec F high* AMs, marked by CD170+, CD11b- and MHC class II-, original *Siglec F low* AMs, marked by CD170 low, CD11b- and MHC class II-, original *Siglec F low,* MHC class II+ AMs marked by CD170 low, CD11b- and MHC class II+, and a new *Siglec F low*, CD11b high AM subset marked by CD170+, CD11b+ and MHC class II-. In contrast to our data, the study showed that the absolute count of original AMs was the same at 14 days post-LPS exposure relative to initial unexposed mice. The absence of AM expansion was further supported by the lack of incorporation of 5-bromo-2′-deoxyuridine (BrdU) in trained macrophages. Although we did not detect expression of the Ki-67 proliferation marker in trained AMs (data are not shown), the absence of Ki-67 does not prevent cells from dividing.^44^ In addition, BrdU staining is undetectable after cells have divided. Thus, rigorous time course BrdU tracking would be required to confirm that the absence of BrdU staining is due to a lack of cell division. Instead, we physically visualized the presence of double-nucleated AMs in the lungs, implying that these cells are undergoing cell division. Nonetheless, our data does not rule out that these cells may also undergo phenotypic plasticity. Thus, we propose that in response to immune stimuli, AMs undergo cell expansion, resilience, and phenotypic plasticity to protect against respiratory pathogens.

Neutrophils are critical components of the protective immune adaptation gained during exposure to immune stimuli; however, there is a double-edged sword to neutrophil activation if their responses are not well controlled.^45,46^ In general, when neutrophils become hyperactivated, they can cause excessive tissue damage that leads to detrimental outcomes. This is especially true if neutrophil hyperactivation occurs in vital organs like the lungs, which often leads to organ failure and death.^47,48^ However, our study infers that increased neutrophil activities can provide a potent protection if they are accompanied by increased clearance of destructive molecules. In agreement with this, several studies demonstrated that exposure to immune stimuli increases neutrophil recruitment and function in response to subsequent challenges, which drives the protective trained immune responses.^35,36,49^ Exposure to LPS or BCG vaccine can induce neutrophils to uniquely express specific genes, which may contribute to neutrophil-mediated killing. Nevertheless, whether exposure to immune stimuli reprograms neutrophils to respond uniquely or induces local microenvironmental changes within organs that modulate neutrophil responses is debatable. Because neutrophils are short-lived cells, it is unlikely that they would be reprogrammed to acquire immunological memory. Instead, several studies have suggested that the ability of neutrophils to acquire immune memory depends on hematopoietic progenitor cells.^50,51^ An initial immune stimuli induces hematopoietic progenitors to undergo metabolic and epigenetic changes, which results in the generation of neutrophils with enhanced antimicrobial functions. In agreement with the previous studies, we found that LPS exposure promotes the release of NETs into BALF to trap and kill MRSA. Importantly, depletion of neutrophils abolished the protection in LPS-exposed mice, which highlights the essential role of neutrophil effector functions. However, adoptive transfer of purified neutrophils from trained mice failed to confer protection in naïve mice. This suggests that changes in the lung microenvironment in response to LPS are still needed for protective neutrophil effector functions. Thus, reprogramed neutrophils or neutrophil-progenitor cells may not be sufficient to provide protection in cellular autonomous manner.

Prior exposure to immune stimuli licenses AMs to acquire neutrophil-derived antimicrobial enzymes from the matrix to kill MRSA and prevent tissue damage. Using primary cell culture, a previous study demonstrated that human macrophages can uptake neutrophil-derived antimicrobial molecules and use them to kill bacteria in phagosomes.^41^ However, mechanisms that control this process and whether it occurs i*n vivo* have remained unknown. Our *ex vivo* microscopy analysis revealed the unanticipated outcome that AMs from LPS-exposed mice are positive for neutrophil-derived enzymes (MPO and Elastase). Although we found that MPO inhibition is insufficient to attenuate the protective response in LPS-exposed mice, other neutrophil defensins could compensate for inhibiting MPO activity. Thus, it remains unanswered whether AMs acquire neutrophil molecules from the extracellular matrix to kill MRSA in phagosomes, prevent tissue damage, or both. However, blocking uptake of MPO by AMs renders LPS-exposed mice highly susceptible to subsequent MRSA challenge. This indicates that the acquisition of neutrophil-derived antimicrobial molecules by AMs is an integral component of innate immune adaptation acquired during initial immune stimulation. Further analysis revealed that several efferocytosis regulators (MerTK, Marco, CD68, and Itgb2) and anti-apoptotic molecules (Bcl-xL and Mcl-1) are specifically induced in LPS-exposed mice. Inhibition of the efferocytosis receptor MerTK or the anti-apoptotic protein Bcl-xL blocks the ability of AMs to uptake neutrophil-derived MPO and renders mice susceptible to MRSA. Interestingly, blocking Bcl-xL did not induce AM cell death, suggesting that it may modulate other cellular functions. In addition to its role as an anti-apoptotic protein, Bcl-xL modulates mitochondrial metabolism,^52–54^ which could drive innate immune adaptations in response to immune stimuli. In summary, capturing neutrophil-released antimicrobial enzymes by AMs to enhance host defenses provides a cooperative host defense strategy between innate immune cells which is an indispensable mechanism of innate immune adaptation acquired during immune stimulation.

## RESOURCE AVAILABILITY

### Lead Contact

All information, including data, materials, and other resources, can be obtained by directing requests to the lead contact, Basel H. Abuaita.

### Materials Availability

All reagents and resources can be shared upon request.

### Data and Code Availability

● Data reported in this manuscript will be shared upon request from the lead contact.
● This paper does not report any new code.
● Any other information needed to reanalyze the reported data is available from the lead contact upon request.

## ACKNOWLEDGMENTS

This work is supported by the LSU Center of Biomedical Research Excellence (COBRE) for lung biology and disease (NIH NIGMS 5P20GM130555; B.H.A). We thank the LSU Bacterial Pathogenesis and Host Response Interest Group members for many helpful discussions. A.S. was supported by Hannelore and Johannes Storz and MegaRobo-Biocytogen Student Travel Awards to attend national conferences.

## AUTHOR CONTRIBUTIONS

A.S. and B.H.A. performed the experiments and wrote the manuscript. L.M.H. and V.M. assisted in experimental preparation. S.D.J. and W.N.B. assisted in designing the experiments and data interpretation.

## STAR Methods

### EXPERIMENTAL MODEL AND SUBJECT DETAILS

#### Mice

Mice used in this study were housed in the Division of Laboratory Animal Medicine (DLAM) at the School of Veterinary Medicine, Louisiana State University (LSU), and A&M College-Baton Rouge. All animal procedures used were approved by the Institutional Animal Care and Use Committee at LSU. Wild-type C57BL/6J, Rag1 KO, and CCR2 KO mice were acquired from Jackson Laboratory. Genotyping was done in-house using the Direct mouse genotyping kit (Cat # K1025, APExBIO) using ear clip samples.

#### Bacteria

A Community-associated Methicillin-resistant *Staphylococcus aureus* (MRSA) strain, USA300 LAC, and its isogenic strain that constitutively expresses GFP (MRSA-GFP) were used in this study.^16,55^ Bacterial stock was stored at -70°C in Luria-Bertani (LB) medium (Cat# 244610, Becton Dickinson) with 20% glycerol. For revival, bacteria were streaked onto tryptic soy medium (Cat# 211825, Becton Dickinson) plates and grown overnight at 37°C. Colonies were randomly selected and cultured overnight in tryptic soy broth (TSB) in a shaking (220 rpm) incubator at 37°C. The next day, bacteria were pelleted, washed, and resuspended in phosphate buffer saline (PBS). The bacterial inoculum was determined based on OD_600_ and confirmed by plating serial dilutions onto the TSB plates to calculate colony-forming units (CFU) present.

#### MRSA-induced pneumonia

Male and female mice at the age of 8-12 weeks were used in the in *vivo* infection experiments. Anesthesia was done by using isoflurane inhalation (3% isoflurane, 2 liters/minute oxygen). The anesthetized mice were placed on a slanted surgical rack and inoculated with either a non-lethal dose (10^6^ bacteria in PBS) or a lethal dose (10^8^ bacteria in PBS) of MRSA via the oropharyngeal route as previously performed.^16^ To model recurrent MRSA infection, mice were initially infected with a non-lethal dose of MRSA for 6 days and subsequently challenged with a lethal dose of MRSA for 48 hours. To model host defense adaptations to immune stimuli, TLR agonists were inoculated to mice oropharyngeal in 50 µl PBS for various periods ranging from day 1 to day 21. LPS (1 µg/mouse) (Cat# ALX-581-008, Enzo Life Sciences) was used to stimulate TLR4 signaling, Pam3CSK4 (50 µg/mouse) (Cat# VAC-PMS, InvivoGen) was used to stimulate TLR2 signaling, and Poly I::C (100 µg/mouse) (Cat# TLRL-PIC, InvivoGen) was used to stimulate TLR3 signaling. Subsequently, mice were challenged with a lethal dose of MRSA. Mice were monitored at least every 6 hours to observe the progression of the infections, mortality, as well as extreme distress in mice. Experimental mice were terminated at 6-, 24-, or 48-hours post-infection to collect bronchoalveolar lavage fluid (BALF) and lungs in PBS. Lungs were further processed by mechanical dissociation to obtain single-cell suspensions as described before.^16,56–58^ The lung single cell suspensions and BALF were used to measure host responses, immune cell recruitments, and bacterial burden by enumeration of CFU. The acellular fractions from single cell suspensions were used to quantify production of inflammatory mediators.

### METHOD DETAILS

#### Quantification of bacterial load and inflammatory mediators

The BALF and lung single-cell suspensions were collected in 3 ml PBS. Samples (50 µl) were first diluted 1:20 in sterile ultrapure water to allow host cell lysis and release of intracellular bacteria. Samples were serially diluted and plated onto TSB agar plates for CFU enumerations. The CFUs were counted using an Acolyte counter and were confirmed by manual counting. Quantification of inflammatory mediators in lungs was done using acellular fractions of lung cell suspensions. First, the lung cell suspensions were centrifuged to pellet cells, and acellular supernatants were collected without disturbing the pellets. Samples were stored immediately at -20°C for future analysis. Multiplex bead-based panels were used to measure the levels of inflammatory mediators simultaneously using flow cytometry, as we previously performed.^16,59^ The LEGENDplex mouse inflammation panel (Cat# 740446, Biolegend) and proinflammatory chemokine panel (Cat# 740451, Biolegend) were used according to the manufacturer’s recommendations. The data was acquired using a BD LSRFortessa flow cytometer and further analyzed using LEGENDplex software suite.

#### Immune cell phenotyping

Immunophenotyping was performed based on specific cell surface markers on cells obtained by centrifugation from BALF and lung single-cell suspensions. The BALF and lung single-cell suspensions were collected from uninfected and infected mice in 3 ml PBS. Samples were centrifuged, and cell pellets were resuspended in red blood cell lysis buffer (Cat# 420301, Biolegend). Cells were collected by centrifugations, washed with PBS, and Fc blocked using TruStain FcX anti-mouse CD16/32 antibody (Cat# 101319, Biolegend). Cells were then surface-stained with a panel of fluorescent conjugated antibodies, including anti-CD45 clone 30-F11, anti-CD11b clone M1/70, anti-CD11c clone N418, anti-CD170 clone S17007L, anti-CD169 clone 3D6.112, anti-Ly-6C clone HK1.4, and anti-Ly-6G clone 1A8 for 30 minutes at room temperature in the dark. Cells were washed twice with staining buffer and analyzed on a BD LSRFortessa flow cytometer. Single staining of UltraComp Plus Compensation Beads (Cat# 01-3333, ThermoFisher) was used for compensation. Flow data was analyzed using FlowJo software. The relative amounts of different immune cells in the BALF and lungs were calculated relative to CD45-positive cells. AMs were identified based on CD11c^+^ and CD170^+^ or CD169^+^, neutrophils were identified based on Ly6G^+^ and CD11b^+^, and monocytes were identified based on Ly6C^+^ and CD11b^+^. Absolute cell counts were calculated using CountBright Absolute Counting Beads for flow cytometry (Cat# C36950, ThermoFisher) according to the manufacturer’s protocol.

#### Cytospin and Diff-Quik staining

BALF cells were cytocentrifuged (4 mins at 2000 rpm) onto slides to immobilize the cells. After overnight drying, slides were differentially stained using the Diff-Quik stain kit (Cat# MER1002, Mercedes Scientific). The staining procedure involved sequential, brief immersion in a methanol-based fixative, Eosin Y solution, a solution with Methylene Blue, and Azure A. Excess stain was rinsed with water, and the slides were dried overnight before taking images on a HAMAMATSU Nanozoomer slide scanner. Image processing was done on OlyVIA software (Olympus).

#### Staining lung histology sections

Mice were euthanized using an automated carbon dioxide delivery system. Immediately following euthanasia, lungs were perfused and inflated by injecting 1.5 ml of 2% low-melting-point agarose in PBS into the mouse trachea. After agarose was solidified, the lungs were excised and fixed in 10% neutral formalin for at least 48 hours. The lobes were separated and embedded in paraffin. Lung sections of 5 µm thickness were cut by the histology core at Louisiana Animal Disease Diagnostic Laboratory (LADDL) and then stained with hematoxylin and eosin (H&E). All H&E slides were then scanned on the HAMAMATSU NanoZoomer Slide Scanner System, and images were further processed using OlyVIA software (Olympus).

#### *Ex vivo* immunofluorescence microscopy

BALF and lung cells were seeded onto polylysine-coated coverslips in 6-well plates. Cells were allowed to settle by gravity and adhere to the coverglass by incubating at room temperature (RT) for 30 minutes. Cells were fixed with Fluorofix buffer (Cat# 422101, Biolegend) for 20 minutes at RT and then stored at 4°C overnight. The next day, cells on coverglass were washed with a wash buffer (0.1% Triton X-100 in PBS) 3 times for a total of 15 minutes to permeabilize the cells. Cells were then blocked by incubating in PBS staining buffer containing 10% normal goat serum (Cat# 10000C, ThermoFisher), 5% Bovine serum albumin (BSA, Cat# BP1600, Fisher Scientific) for 30 minutes. Cells were stained for 45 minutes at RT with primary antibodies in staining buffer. The following antibodies were used: anti-CD169 clone 3D6.112 (Biolegend), anti-CD11C clone D1V9Y (Cell signaling), anti-MPO polyclonal (Dako), anti-Lamp1 clone 1D4B (DSHB), and anti-neutrophil elastase (NE) polyclonal (Abcam). Cells were washed 3 times with a wash buffer and then incubated for 45 minutes at RT in the dark with fluorescent-conjugated Alexa Fluor secondary antibodies (ThermoFisher). Cells were counterstained using DAPI for 10 minutes and washed at least 3 times for a total of 15 minutes. Coverslips were then mounted onto a glass slide using ProLong Diamond Antifade Mountant (Cat# P36961, ThermoFisher). Cells were imaged using an Olympus FV3000 confocal microscope, and MFIs were quantified using ImageJ software.

#### Real-time quantitative PCR (qPCR) and NanoString assay

RNA samples were isolated from lung cells using the mirVana miRNA isolation kit (Cat# AM1561, ThermoFisher). RNA samples were quantified by NanoDrop spectrophotometer and normalized across all conditions. cDNA samples were synthesized using random hexamers (Cat# N8080127, ThermoFisher), dNTP Mix (Cat# 18427088, ThermoFisher), M-MLV Reverse Transcriptase (Cat# 28025013, ThermoFisher), and RNase Inhibitor (Cat# N8080119, Applied Biosystems). The qPCR was performed using Applied Biosystems PowerUp SYBR Green Master Mix (Cat# A25778, ThermoFisher) with following parameters: [cycle 1: 50°C for 2 mins, cycle 2: 95°C for 2 mins, cycle 3: 95°C for 15 secs, 55°C for 15 secs, 72°C for 1 min (40 repeats), cycle 4: 95°C for 15 secs, 60°C for 15 secs and 95°C 15 secs]. Expression levels were normalized using *Gapdh* and calculated as fold change (2^-ΔΔCt^) relative to the uninfected control group. Primer sequences used in this study are listed in our previous study.^16^ RNA samples from lungs of uninfected mice (PBS), MRSA-infected naive mice, and MRSA-infected mice after exposure to LPS for 6 days were subjected to NanoString analysis to quantify expression of inflammatory genes. The nCounter mouse host response panel and the nSolver 4.0 software (NanoString Technologies) were used to quantify expression of 785 inflammatory genes.

#### Innate immune cell depletions

AMs were depleted using Clodronate-coated liposomes (Cat# CP-005-005, Liposoma BV). Control or Clodronate liposomes were inoculated in 50 μl PBS via oropharyngeal aspiration over two consecutive days. AM depletions were confirmed by quantifying the absolute AM counts in BALF samples. To deplete blood innate immune cells, mice were injected with 0.2 mg in 200 μl of rat IgG2α isotype control (Cat# BE0089, BioXCell), anti-mouse Ly6G monoclonal antibody for neutrophil depletion (Cat# BE0075-1, BioXCell), or anti-mouse Ly6C monoclonal antibody for monocyte depletion (Cat# BE0203, BioXCell). Injections were performed intravenously through the tail vein for two consecutive days. Depletion efficiencies were calculated by the percent reduction of the corresponding cell type in MRSA-infected lungs.

#### *In vivo* rhodamine liposome cell labeling

Rhodamine packaged liposomes were purchased from Liposoma BV and used to label AMs or blood monocytes. Rhodamine is a stable molecule with bright red fluorescence that was previously used to label phagocytes after being packaged into liposomes.^60^ To label AMs, mice were inoculated with 100 µl of rhodamine liposomes via the oropharyngeal route. BALF cells were collected 24 hours after inoculation and visualized by microscopy to confirm uptake of the rhodamine liposomes by AMs. To label blood monocytes, mice were injected with 200 µl of rhodamine liposomes intravenously through the tail vein. Cells from whole blood were collected 24 hours after injection, and the efficiency of monocyte labeling was quantified by flow cytometry using monocyte markers (anti-Ly6C and anti-CD11b). To trace labeled cells, rhodamin liposomes-inoculated mice were left to recover for 24 hours before LPS administration via the oropharyngeal route for 6 days. Subsequently, mice were challenged with the MRSA-GFP challenge via the oropharyngeal route for 48 hours. BALF cells were collected, seeded onto polylysine-coated coverslips, and stained with anti-CD169 clone 3D6.112 and DAPI. Cells were imaged using an Olympus VF3000 confocal microscope and processed using cellSENS imaging software.

#### Neutrophil purifications and adoptive transfer

Bone marrow cells were extracted from naive mice or LPS-exposed mice, oropharyngeally, for 6 days. Bone marrow from mouse femurs and tibias was flushed with PBS, and cells were mechanically dissociated using a syringe handle on a 70 µm nylon mesh. The Mojosort mouse neutrophil extraction kit (Cat# 480058, Biolegend) was used to purify neutrophils using the manufacturer’s protocol. Neutrophil purity was confirmed by flow cytometry analysis of cells stained with neutrophil markers (anti-Ly6G and anti-CD11b). Purified neutrophils were resuspended in PBS, normalized, and adoptively transferred to naïve mice via tail vein injection. Each mouse was administered 10^6^ cells in 200 µl of PBS. Mice were challenged with a lethal dose of MRSA after 30 minutes of neutrophil adoptive transfer via oropharyngeal inoculation.

#### *In vivo* pharmacological inhibition

All inhibitors were purchased from MedChemExpress (MCE), resuspended in sterile DMSO at 50 mg/ml stocks, and stored at -20°C. Venetoclax was used to inhibit Bcl-2 (Cat# HY-15531, MCE), UNC2025 was used to inhibit MerTK (Cat# HY-12344, MCE), and A-1331852 was used to inhibit Bcl-XL (Cat# HY-19741, MCE). Inhibitors (50 mg/ml stock) or control DMSO were diluted 1:10 in PBS and used to treat mice with 50 µl aspiration via the oropharyngeal route. The next day, mice were infected with MRSA along with inhibitors. The concentration of inhibitors used did not interfere with MRSA survival or growth, which was confirmed by the enumeration of similar bacterial counts in all inoculations.

#### Live confocal microscopy

BALF cells were extracted from naive or LPS-exposed mice in 2 ml PBS after MRSA infection (24 hpi). Cell suspensions were layered onto Histopaque1119 density gradient (Cat# 11191, MilliporeSigma) and centrifuged at 1500 g for 30 min to separate RBC from leukocytes. Leukocyte layers were collected, diluted with PBS, and adhered onto a polylysine-coated glass-bottom dish for 20 mins at RT. Cells were washed with PBS and cultured in RPMI media without phenol red and with 10% FBS. Cells were stained with Hoechst 33342 DNA dye (Cat# 62249, ThermoFisher) and imaged live immediately using an Olympus FV3000 confocal microscope. Images were further processed using cellSENS imaging software.

#### Statistical analysis

All *in vivo* experiments were done at least two times on different days. Graphs represent mean ± standard deviation or mean ± standard error of mean as indicated in figure legends. GraphPad Prism 10 was used for graphing and performing statistical analysis. The Mann-Whitney test was used to determine significant differences between two groups. One-way ANOVA with Holm-Sidak’s post-test was used to determine significant differences between three or more groups. Two-way ANOVA followed by Holm-Sidak’s multiple comparisons test was used to determine significant differences between groups that were defined by columns and rows. Log-rank (Mantel-Cox) test was used to calculate p-values of survival curves. All p-values < 0.05 were considered significant, and all statistically significant comparisons in groups were indicated. \**P* < 0.05; \*\**P* < 0.01, \*\*\**P* < 0.001, \*\*\*\**P* < 0.0001, ns: not statistically significant.

**Supplemental Figure 1.**
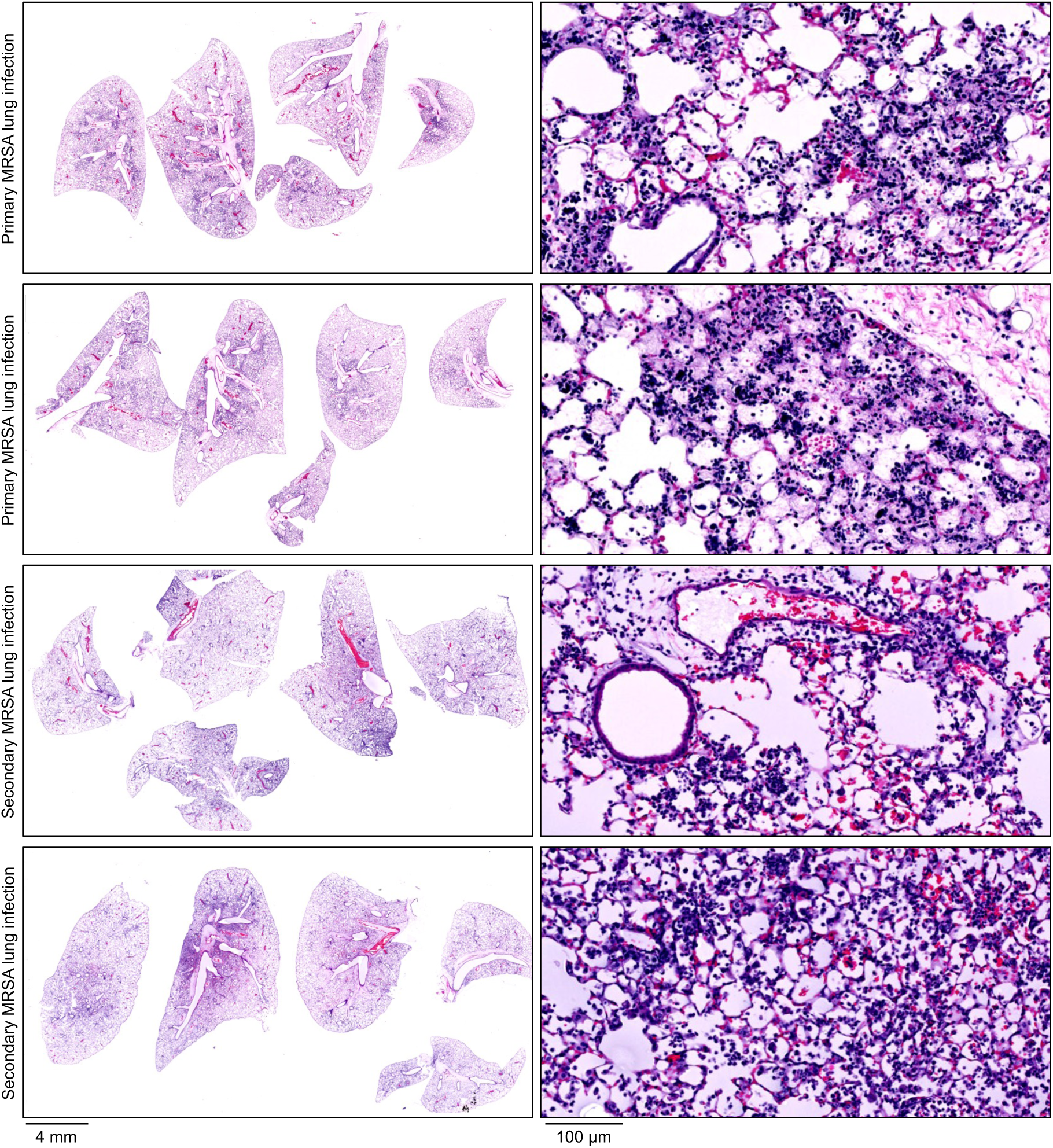
Exposure to non-lethal MRSA reduces lung injuries during subsequent lethal MRSA infections. Hematoxylin and eosin (H&E) stained lung histology sections from primary or secondary MRSA-infected mice. Mice were inoculated with phosphate buffer saline (PBS) or a non-lethal dose (10^6^ CFU) of MRSA for 6 days and then challenged with a lethal dose (10^8^ CFU) of MRSA for 48 hours. All inoculations were performed by oropharyngeal aspiration.

**Supplemental Figure 2.**
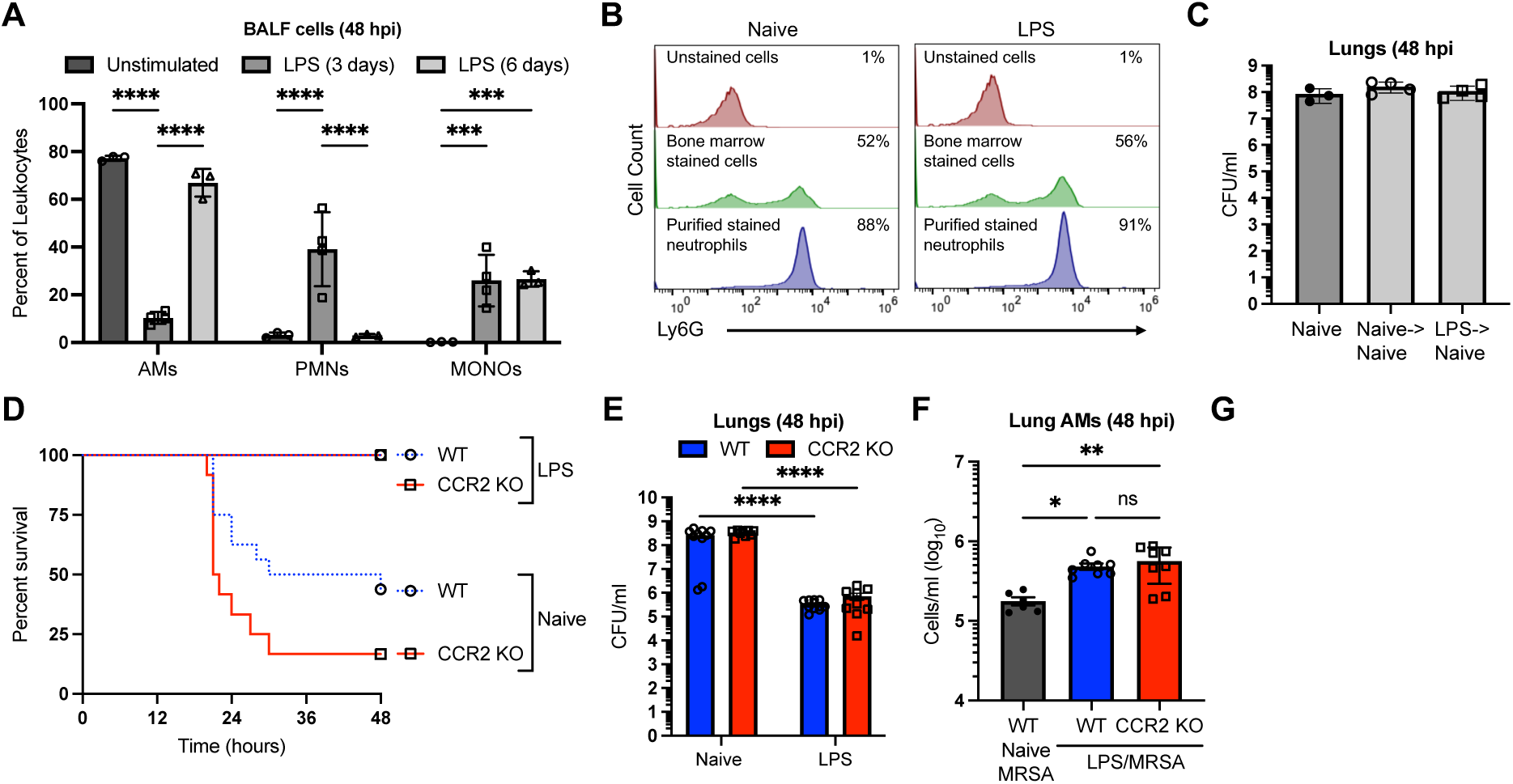
Neutrophils and CCR2-mediated inflammatory monocyte recruitments are insufficient for the LPS-induced immune protection against MRSA infections. (A) Immunophenotyping analysis of leukocytes in BALF of naive mice (n=4) and mice exposed to LPS via oropharyngeal aspiration for 3 or 6 days (n=4). Cells were stained for AMs using anti-CD11c, anti-CD170, neutrophils (PMNs) using anti-Ly6G, anti-CD11b, and monocytes (MONOs) using anti-Ly6C, anti-CD11b, and analyzed by flow cytometry. (B) Flow cytometric analysis of the percentage of neutrophils in bone marrow and its neutrophil-purified fraction from naive and LPS-exposed mice. Cells were stained with anti-CD45, anti-Ly6G, and anti-CD11b. The percent of neutrophils was determined by gating against unstained cells. (C) Bacterial burden in lungs of mice when injected with PBS (naive), adoptively transferred with purified neutrophils from naive mice (naive->naive), or adoptively transferred with purified neutrophils from LPS-exposed mice (LPS->naive) one hour prior to MRSA challenges. (D) Survival curves of MRSA-infected WT (n≥8) and CCR2 KO (n≥8) mice when mice were injected with PBS (naive) or LPS via oropharyngeal aspiration for 6 days prior to infections. (E) Bacterial burden in lungs of MRSA-infected naive WT mice (n=10), naive CCR2 KO mice (n=9), LPS-exposed WT mice (n=11), and LPS-exposed CCR2 KO mice (n=9). (F, G) Alveolar macrophage count (F) and monocyte counts (G) in lungs of naive WT mice (n=6), LPS-exposed WT mice (n=8), and LPS-exposed CCR2 KO mice (n=8). Graphs representing the mean from at least two independent experiments ± SEM. Statistical analysis was performed by Two-way ANOVA for panels A and E with Holm-Sidak’s post-test for multiple comparisons and One-way ANOVA with Holm-Sidak’s post-test for panels C, F, and G. *P* value: *<0.05; **< 0.01, ***< 0.001, ****< 0.0001, and ns: not statistically significant.

**Supplemental Figure 3.**
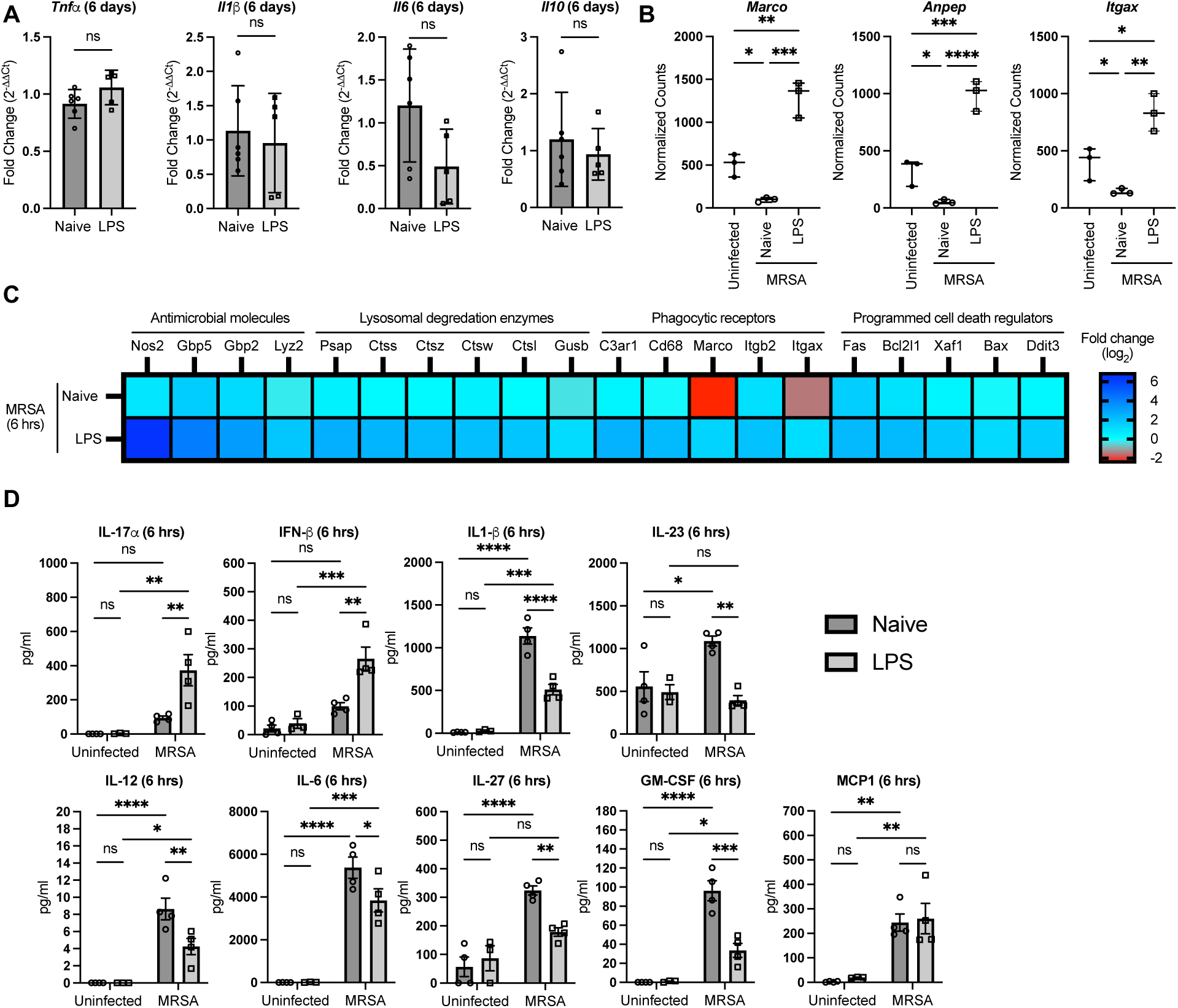
Expression of inflammatory genes is differentially changed in lungs of LPS-exposed mice relative to unexposed mice during MRSA infections. (A) qPCR of *Tnfα*, *Il1β*, *Il6,* and *Il10* transcript levels in lungs of uninfected mice when mice were exposed to LPS for 6 days or left untreated (naive). (B) Transcript levels of *Marco*, *Anpep*, and *Itgax* in lungs of uninfected mice (n=3), MRSA-infected mice pre-exposed to LPS for 6 days (n=3), or MRSA-infected naive mice (n=3). Graphs represent the mean of normalized counts ± SD from the NanoString data. (C) Expression of selected genes from the NanoString data presented as log_2_(fold change) in lungs of MRSA-infected naive mice or mice exposed to LPS for 6 days relative to lungs from uninfected mice. (D) Levels of indicated inflammatory mediator in supernatants of lung signal-cell suspensions from uninfected naive mice, uninfected LPS pre-exposed mice, MRSA-infected naive mice, and MRSA-infected LPS pre-exposed mice. Graphs represent the mean of n≥3 biological replicates ± SD. *P* values were calculated using the Mann-Whitney test for panel A, One-way ANOVA followed by a Holm-Sidak’s multiple comparisons test for panel B, or Two-way ANOVA followed by a Holm-Sidak’s multiple comparisons test for panel D. *P* value: *<0.05; **< 0.01, ***< 0.001, ****< 0.0001, and ns: not statistically significant.

**Supplemental Figure 4.**
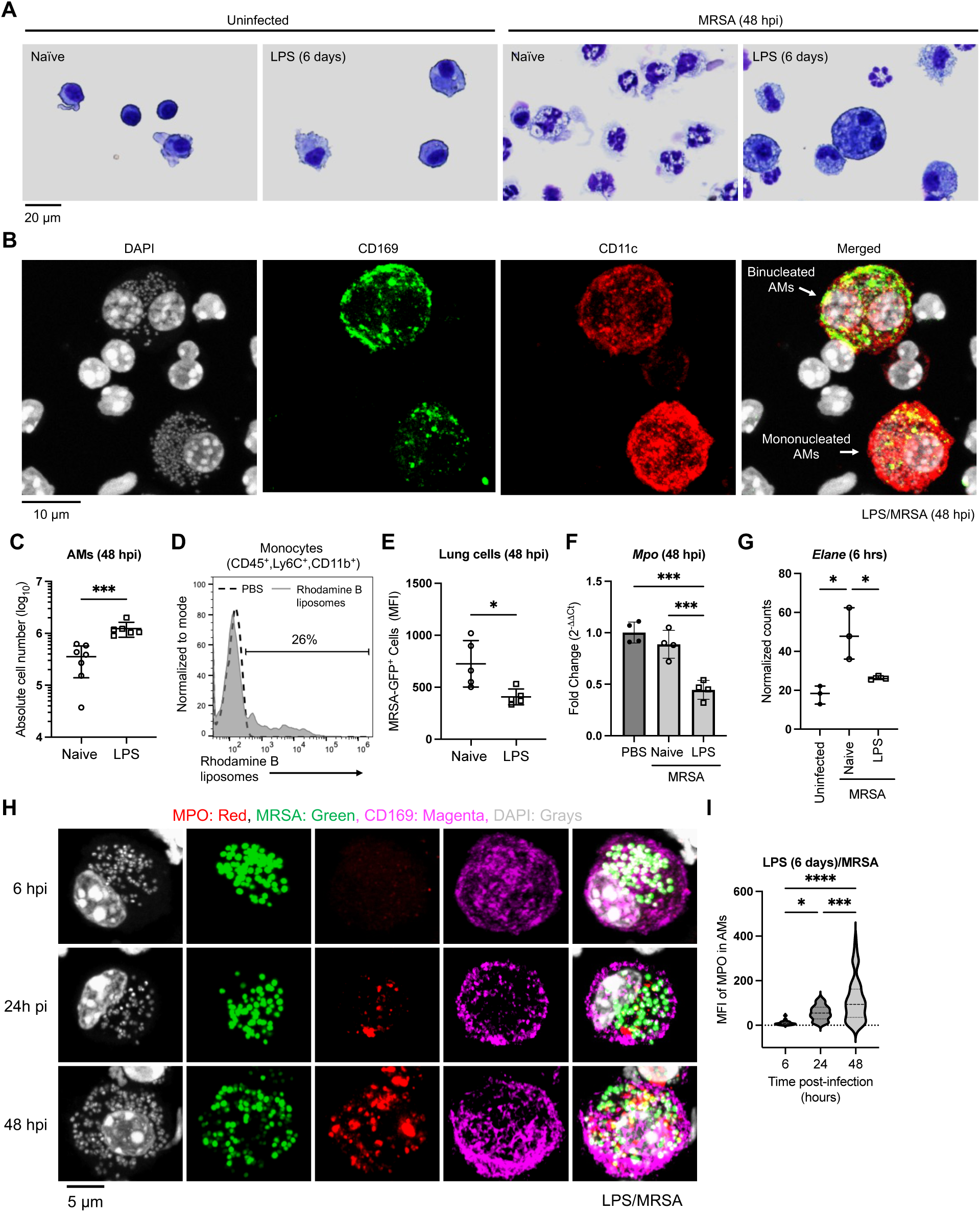
LPS exposure stimulates AM self-renewal and MPO uptake. (A) Representative cytospin image of BALF cells from uninfected naive mice, uninfected LPS pre-exposed mice, MRSA-infected naive mice, or MRSA-infected LPS pre-exposed mice. Scale bar represents 20 µm. (B) Representative confocal microscopy images or Mono and Binucleated AMs from LPS pre-exposed mice for 6 days and subsequently infected with MRSA for 48 hours. Cells were extracted from the lungs ex vivo and stained with DAPI (gray), CD169 (green), and CD11c (red). Scale bar represents 10 µm. (B) Absolute AM counts in lungs of MRSA-infected mice when mice were left untreated (naive) or pre-exposed to LPS for 6 days. (D) Representative flow cytometry histogram of blood monocytes from mice injected with control PBS or Rhodamine liposomes via the tail vein. Cells were stained with monocyte markers (CD45, CD11b, and Ly6C). The percent of monocyte-positive liposomes was determined by gating against monocytes from PBS-injected mice. (E) Mean fluorescence intensity of GFP in lung cells when naive or LPS pre-exposed mice infected with MRSA-GFP for 48 hours. (F) The expression level of *Mpo* was quantified by qPCR and normalized relative to *Gapdh*. The ΔΔCt was calculated relative to the average ΔCt of PBS-injected (uninfected) mice. (G) Quantification of *Elane* transcripts in lungs of uninfected mice (n=3), MRSA-infected naive mice (n=3), and MRSA-infected LPS pre-exposed mice (n=3) from NanoString data. (H, I) Representative confocal microscopy images (H) and quantification (I) of MPO levels in AMs of LPS pre-exposed mice when mice were infected with MRSA-GFP for 6, 24, or 48 hours. Scale bar represents 5 µm. Graphs represent the mean of n≥3 biological replicates ± SD. *P* values were calculated using the Mann-Whitney test for panels C and E, or One-way ANOVA followed by a Holm-Sidak’s multiple comparisons test for panels F, G, and I. *P* value: *<0.05; **< 0.01, ***< 0.001, ****< 0.0001.

**Supplemental Figure 5.**
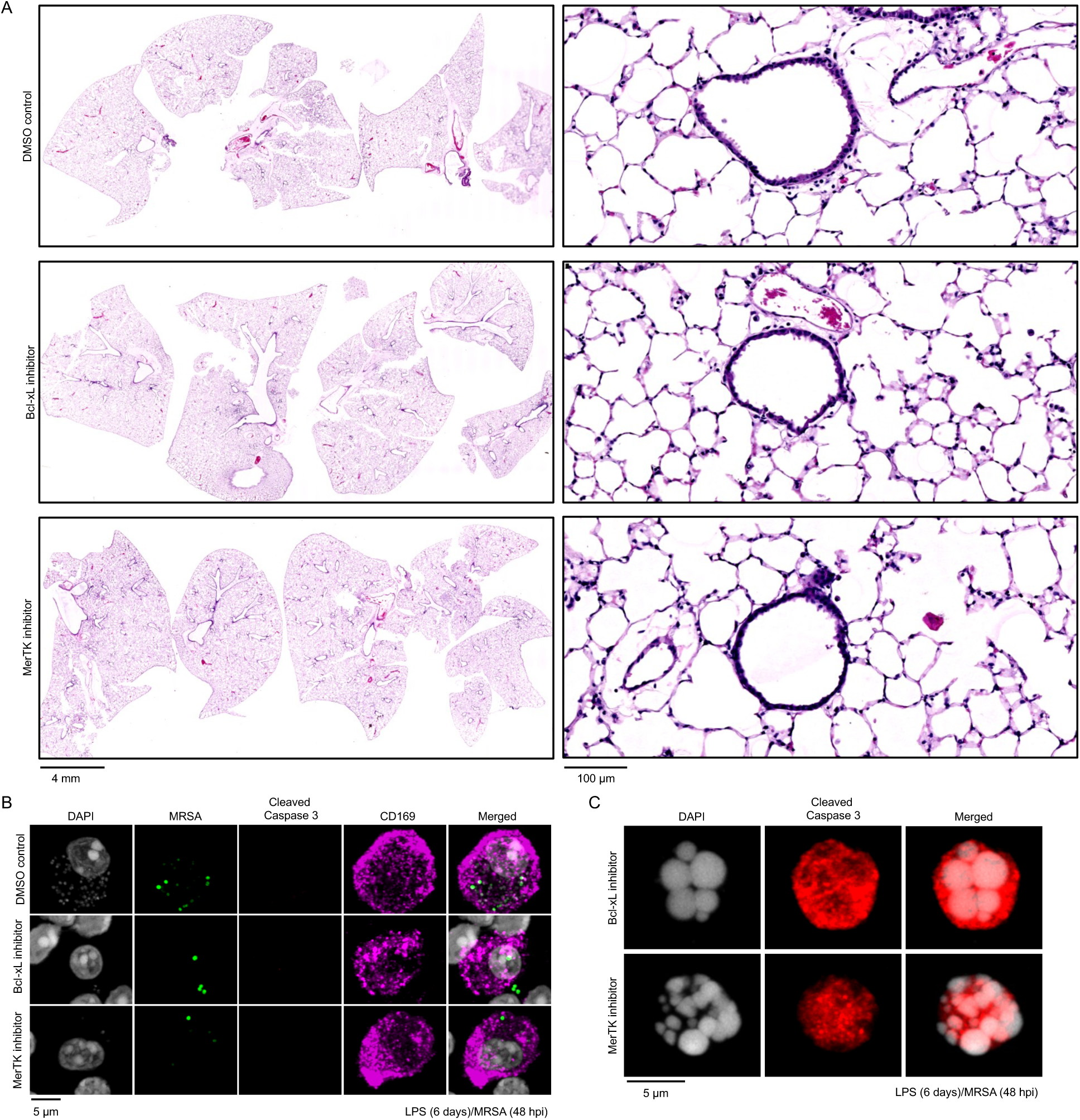
Bcl-xL and MerTK inhibitors do not cause lung toxicity to uninfected mice or MRSA-infected AM cell death in LPS-exposed mice. (A) H&E-stained lung histological sections from mice treated with DMSO control, Bcl-xL inhibitor (A-1331852), or MerTK inhibitor (UNC2025). (B) Representative confocal microscopy images of AMs extracted from lungs of LPS-exposed mice for 6 days, when mice were treated with DMSO control or Bcl-xL inhibitor, or MerTK inhibitor before being infected with MRSA-GFP (green) for 48 hours. Cells were stained with DAPI (gray), anti-cleaved Caspase 3 (red), and anti-CD169 (magenta) Scale bar represents 5 µm. (C) Representative confocal microscopy images of cleaved Caspase 3-positive cells present in lungs of LPS-exposed mice, treated with Bcl-xL or MerTK inhibitors, and infected with MRSA. Scale bar represents 5 µm.

